# Identification of a dominant chlorosis phenotype through a forward screen of the *Triticum turgidum* cv. Kronos TILLING population

**DOI:** 10.1101/622076

**Authors:** Sophie A. Harrington, Nicolas Cobo, Miroslava Karafiátová, Jaroslav Doležel, Philippa Borrill, Cristobal Uauy

## Abstract

Durum wheat *(Triticum turgidum)* derives from a hybridization event approximately 400,000 years ago which led to the creation of an allotetraploid genome. Unlike with more ancient whole genome duplications, the evolutionary recent origin of durum wheat means that its genome has not yet been fully diploidised. As a result, many of the genes present in the durum genome act in a redundant fashion, meaning that, in many cases, loss-of-function mutations must be present in both gene copies to observe a phenotypic effect. This redundancy has hindered the use of forward genetic screens in durum wheat. Here we use a novel set of induced variation within the cv. Kronos TILLING population to identify a locus controlling a dominant, environmentally-dependent chlorosis phenotype. We carried out a forward screen of the sequenced cv. Kronos TILLING lines for senescence phenotypes and identified a single line with a dominant early senescence and chlorosis phenotype. Mutant plants contained overall less chlorophyll throughout their development and displayed premature flag leaf senescence. A segregating population was classified into discrete phenotypic groups and subjected to bulked-segregant analysis using exome capture followed by next-generation sequencing. This allowed the identification of a single region on chromosome 3A, *Yellow Early Senescence 1 (YES-1),* which was associated with the mutant phenotype. To obtain further SNPs for fine-mapping, we isolated chromosome 3A using flow sorting and sequenced the entire chromosome. By mapping these reads against both the cv. Chinese Spring reference sequence and the cv. Kronos assembly, we could identify high-quality, novel EMS-induced SNPs in non-coding regions within *YES-1* that were previously missed in the exome capture data. This allowed us to fine-map *YES-1* to 4.3 Mb, containing 59 genes. Our study shows that populations containing induced variation can be sources of novel dominant variation in polyploid crop species, highlighting their importance in future genetic screens. We also demonstrate the value of using cultivar-specific genome assemblies alongside the gold-standard reference genomes particularly when working with non-coding regions of the genome. Further fine-mapping of the *YES-1* locus will be needed to identify the causal SNP underpinning this dominant, environmentally dependent phenotype.

## Introduction

Polyploidisation events underpin plant evolution and have been suggested to be key drivers of innovation, particularly within the angiosperms (Soltis and Soltis, 2016). All angiosperm species, including important crops such as wheat, rice, and maize, carry signatures within their genomes of ancient whole genome duplication (WGD) events that occurred within their lineage, such as the monocot-specific duplication, τ (Paterson et al., 2012). These Polyploidisation events lead to the presence of multiple copies of genes which previously carried out the same function. It has been proposed that, following WGD, the resulting diploidisation of the genome leads to neo-functionalization or sub-functionalisation of gene copies derived from the original WGD (Dodsworth et al., 2016, Clark and Donoghue, 2018). The diploidisation process reduces the redundancy present within the genome by minimising the number of genes with duplicate functions.

However, unlike rice and maize, wheat has also undergone two more recent allopolyploidisation events, where inter-species hybridizations bring together the chromosomes of each parent, creating a hybrid species with higher ploidy. The first event, approximately 400,000 years ago, occurred when two wild grasses hybridized to produce a tetraploid grass (wild emmer) which would go on to be domesticated as pasta, or durum, wheat *(Triticum turgidum)* (Dubcovsky and Dvorak, 2007, Borrill et al., 2019). The second Polyploidisation event occurred more recently, only 10,000 years ago, when the tetraploid emmer hybridised with another diploid wild grass, leading to a hexaploid species which was then domesticated as bread wheat *(Triticum aestivum).* Unlike in ancient WGDs, these polyploidisation events have occurred relatively recently, such that most wheat genes are present as homoeologous duos or triads in pasta and bread wheat, respectively, and may often have redundant functions (Ramírez-González et al., 2018).

A direct result of this homoeolog redundancy is that the inheritance of many traits in polyploid wheat tend to be quantitative, with multiple homoeologous loci contributing partly to the phenotype (Borrill et al., 2019, Brinton and Uauy, 2019). The phenotypic consequences of mutations in single homoeologs in wheat can be broadly classified into three categories — dominant (e.g. *VRN1),* whereby the mutant allele leads to a complete change in phenotype akin to mutations in diploids (Yan et al., 2003); additive (e.g. *NAM, GW2),* whereby mutants in each homoeolog lead to a partial change in phenotype which becomes additive as mutations are combined (Avni et al., 2014, Pearce et al., 2014, Borrill et al., 2018, Wang et al., 2018); and full redundancy (e.g. *MLO),* whereby the single and double mutants are similar to wildtype individuals, and only the full triple mutant leads to significant phenotypic variation (Acevedo-Garcia et al., 2017). The presence of homoeolog redundancy, therefore, can hinder the use of forward genetic screens in polyploid wheat.

Therefore, beyond its status as an important crop, tetraploid pasta wheat can provide a useful system to reduce the redundancy inherent in polyploid wheat. New advances in wheat genomics resources are increasing the speed and resolution with which we can now map loci corresponding to quantitative traits (Uauy, 2017). Recently gold-standard reference genomes for wheat were released, based on the hexaploid landrace Chinese Spring (IWGSC et al., 2018) and the tetraploid cultivar Svevo (Maccaferri et al., 2019). Additional wheat cultivars from across the globe are being sequenced as part of the wheat 10+ pan-genome project (10+ Wheat Genomes Project, 2016). Crucially, this also includes durum wheat cultivar Kronos, which was used in the development of an *in silico* TILLING population (Krasileva et al., 2017). This mutant resource contains over 4M chemically-induced point mutation variation that can be rapidly accessed for a gene of interest through Ensembl Plants (Vullo et al., 2017).

An additional challenge when working in wheat is the sheer size of the genome, approximately 16 Gb in hexaploid and 11 Gb in tetraploid wheat. This is particularly important when designing sequencing strategies of mutant populations or individuals for mapping-by-sequencing. Various reduced representation methods exist for subsampling the wheat genome. These include gene-based methods through exome capture (Mamanova et al., 2010, Krasileva et al., 2017) or sequencing a specific gene family, as in R-gene enrichment sequencing (RenSeq) (Jupe et al., 2013, Steuernagel et al., 2016). However, these methods are less successful in obtaining variant information from non-coding regions due to their focus on genic regions. This is particularly important in the case of dominant phenotypes, which are often due to variations in regulatory regions that are not within the gene body (Yan et al., 2004, Fu et al., 2005, Borrill et al., 2015), although not exclusively (Simons et al., 2006, Greenwood et al., 2017). Methods do exist, however, to facilitate subsampling of the wheat genome while still retaining information from non-coding regions. In particular, chromosome flow sorting reduces the size of the genome by isolating an entire chromosome which can then be sequenced (Doležel et al., 2012). Other techniques (implemented in rice) include skim sequencing, which uses low coverage to obtain information about deletions or duplications, as well as SNPs, across the genome (Huang et al., 2009).

Here we use the Kronos TILLING population as a case study to identify and fine-map a novel locus in a tetraploid background (Krasileva et al., 2017). We performed a forward screen of the Kronos TILLING population for lines that exhibited late or early senescence phenotypes. From this set, we identified a line that segregated for a dominant chlorosis phenotype and was consistent across multiple years of field trials. We used mapping-by-sequencing to define the dominant phenotype as a single Mendelian locus on chromosome 3A, which we called *Yellow Early Senescence-1*. Using exome capture and chromosome flow-sorting to subsample the large wheat genome, we utilised the new RefSeqv1.0 hexaploid reference genome (IWGSC et al., 2018) alongside an assembly of the durum cultivar Kronos to identify SNPs across the region of interest. Following this, we mapped the *Yellow Early Senescence-1* locus to 4.3 Mb, containing 59 high-confidence genes.

## Methods

### Field Trials

#### TILLING population screen

The initial screen of the sequenced Kronos TILLING population (N=951 M_4_ lines) was carried out on un-replicated single 1 m rows (Supplementary Figure 1A), sown in November 2015 at Church Farm, Bawburgh (52°38’N 1°10’E). Note that all John Innes Centre (JIC) trials were sown at Church Farm, but in different fields at the farm in each year. Lines were sown in numerical order (i.e. line Kronos0423 was followed by Kronos0427). For simplicity, TILLING lines will be referred to as KXXXX throughout the manuscript (i.e. Kronos0423 as K0423). Wild-type controls (cvs. Kronos, Paragon, and Soissons) were sown randomly throughout the population. Rows were phenotyped for senescence as detailed below. Following scoring, 10 mutant lines with early flag leaf and/or peduncle senescence and 11 mutant lines with late flag leaf and/or peduncle senescence were crossed in the glasshouse to wild-type Kronos (Supplementary Table 1). The F_1_ plants were then self-pollinated to obtain F_2_ seed (Fig. 1E). For three mutant lines (K0331, K3085 and K3117) we recovered insufficient F_2_ seeds and hence these populations were not pursued further. All original mutant lines described are available through the JIC Germplasm Resources Unit (www.seedstor.ac.uk).

**Figure 1:**
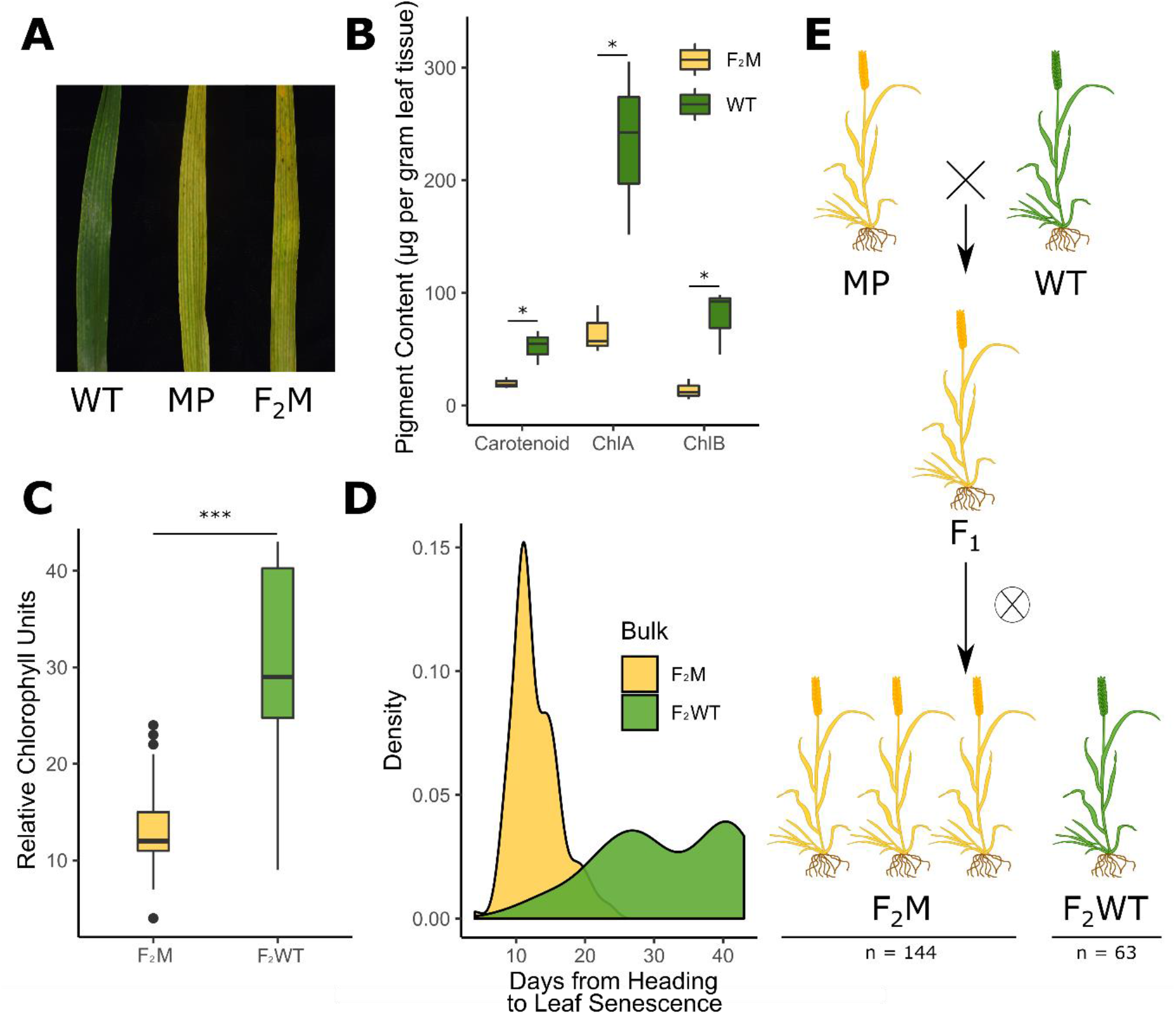
A premature yellowing phenotype from the Kronos TILLING population segregates as a single dominant locus. F_2_ populations of the K2282 Kronos TILLING line grown at the JIC in 2016 showed an early yellowing phenotype (A). Pigment content was measured in the yellow mutant plants (F_2_M) compared to the wild-type plants (F_2_WT) (B; n = 3 per genotype) and was also quantified using SPAD (C; n = 153 F_2_M, n = 61 F_2_WT). The yellow group (F_2_M) senesced significantly earlier than the late bulk (F_2_WT) (D; n = 148 F_2_M, n = 56 F_2_WT). Scoring of the plants demonstrated that the F_2_ population was segregating 3:1 for the yellow trait, indicative of a dominant single locus (E; numbers are combined for both populations). F_2_M and F_2_WT refer to plants which are yellow and green, respectively, and which derive from the F_2_ population (see bottom of E), while WT and MP refer to Kronos WT plants or M_4_ K2282 plants, respectively (see top of E).

#### Recombinant Scoring

F_2_ populations of the selected TILLING lines (backcrossed to cv. Kronos) were sown at Church Farm in March 2016 and grown as described previously (Harrington et al., 2019). Briefly, individual F_2_ seeds were hand-sown in 6×6 1 m^2^ grids, leaving approximately 17 cm between each plant (Supplementary Figure 1B). In total, we sowed 31 F_2_ populations representing 18 distinct TILLING mutant lines. For K2282, two F_2_ populations were sown, K2282-28 and K2282-23, and phenotyped. Seeds from both K2282 populations were taken forward for further field trials.

In 2017 and 2018, the F_3_ seed from the K2282 F_2_ plants that were either heterozygous across the identified region on chromosome 3A or contained recombination within the mapped interval were grown. In 2017, we selected 30 lines from the K2282-28 population and 8 lines from the K2282-23 population. F_3_ seed from these 38 lines were sown in a randomized block design, replicated between 1 to 4 times depending on seed availability. Each experimental unit consisted of a 1 m^2^ plot that contained three 1m rows of a single lines, separated from each other by ~17 cm (Supplementary Figure 1A). The primary tillers of 12 individual plants from each row were tagged before heading. In 2018, 374 individual seeds derived from 16 F_2_ plants completely heterozygous across the SH467/SH969 region were hand-sown into a 1 m^2^ grid (Supplementary Figure 1B) and scored as in 2016. In each year, tissue for genotyping was sampled from the tagged plants (2017) or each individual plant (2018). Senescence phenotyping was carried out as detailed below. Precipitation data for the JIC field trials was obtained from a weather station at 52°37’52.29” N, 1°10’23.57” E.

#### Phenotypic Characterisation

Based on the 2017 genotypic information, nine individual F_3_ lines genotyped as fully homozygous mutant (N = 4) or homozygous wild-type (N = 5) across the initial mapped region (from marker SH467 to SH969) were selected. Plants from these nine genotypes were sown in 2018 in 1 m^2^ plots (two double 1 m rows separated by approximately 33 cm; Supplementary Figure 1C) and replicated 3 times in a complete randomized design. Wildtype Kronos and M5 seed from the K2282 line were also sown as controls (N = 3). Two tillers in each row were tagged at heading and used for SPAD readings and genotyping. Senescence phenotyping was carried out as detailed below.

#### Davis, California Trial

91 F_3_ lines from population K2282-28 and 55 F_3_ lines from population K2282-23 were sown at the University of California Field Station near Davis, California (38° 31’ N, 121° 46’ W) in November 2016. Lines were selected if the F_2_ parent contained recombination within the SH467/SH969 region or was fully heterozygous across the region. In addition, seed from F_2_ parents completely mutant or wild-type across the region (N = 12 each) were also selected. Lines were sown in a complete randomised design, as double 1m rows each separated by an empty row (as in the JIC 2018 trial; Supplementary Figure 1C). Eight individual plants were tagged per row at heading for plants derived from heterozygous parents, to allow genotyping and scoring of individual plants. At least two plants per double row were also tagged and sampled to verify the genotype of the completely wild-type or mutant lines. Heading and visual senescence was scored as in 2018 at the JIC, detailed below. 200 lb N/acre was applied to the trial (as ammonium sulphate), half before planting and half on March 31^st^ (Z30 stage). The trial was treated with appropriate fungicides to prevent stripe rust *(Puccinia striiformis* f. sp. *tritici).* Precipitation data for the Davis trial was obtained from the Davis, California weather station (38° 32’ 07’’ N, 121° 46’ 30’’ W).

#### Glasshouse Trial

F_3_ plants derived from mutant or wild-type F_2_ parents, genotyped across the SH179-SH969 region, were pre-germinated on damp filter paper for 48 hrs at 4°C in the dark. The seedlings were sown into P96 trays with 85% fine peat and 15% horticultural grit. Plants were transplanted to 1L pots at the 3-leaf stage. The pots contained either a) Petersfield Cereal Mix (Petersfield, Leicester, UK), b) Horticultural Sand (J. Arthur Bower’s, Westland Horticulture), or c) Soil taken from the Church Farm site used for JIC field trials (Bawburgh, UK). Plants sown into sand were also supplied with 100 mL of Hoagland solution every three days (Hoagland and Arnon, 1950). K2282 mutant and wild-type F_3_ plants were also tested under low water conditions in each of the three soil conditions listed above. Under the low water conditions, the plants were watered once weekly, and additionally to maintain a soil volumetric water content of approximately 20%, as measured with the Decagon GS3 sensor (ICT International, Armidale, Australia). Three plants of each genotype were treated in each condition. Plants were visually phenotyped for chlorosis onset, determined as a visual yellowing of the main flag leaf (see Figure 1A for a visual example).

### Plant Phenotyping

#### Senescence Phenotyping

Plants were scored for senescence across the different field trials as detailed previously (Harrington et al., 2019). Briefly, when scoring individual plants, all phenotyping was carried out on the main tiller, tagged upon heading. Heading was scored at Zadoks growth stage 57, when the spike was 25% emerged (Zadoks et al., 1974). Flag leaf senescence was scored for the main tiller when 25% of the flag leaf showed visual yellowing and tissue death (necrosis) from the tip. Senescence of the main peduncle was scored when the top inch was fully yellow. When scoring rows of the same genotype, all stages were scored across the entirety of the row. Rows were considered to have reached heading when 75% of the main spikes reached Zadoks growth stage 57. Leaf senescence was similarly scored when 75% of the flag leaves were yellowing and necrotic across 25% of the leaf, from the tip. Peduncle senescence was scored when the top inch of 75% of the peduncles were completely yellow.

Alongside visual scoring, we utilised the SPAD-502 meter (Konica Minolta, Osaka, Japan) to obtain non-destructive chlorophyll content readings. For measurements of individual plants (2016, 2017, 2018) eight readings were taken along the flag leaf on each side of the midrib and averaged to obtain a final reading which was considered the SPAD score for that biological replicate. For measurements of rows (2018), the two tagged tillers were both measured in the same way, and the average of their measurements was taken as the SPAD reading for that biological replicate.

#### Chlorophyll Quantification

Chlorophyll content was measured directly from sampled leaf tissue in 2016 and 2018 at JIC. In 2016, flag leaf tissue was sampled at heading (N = 3 per genotype); in 2018 flag leaf tissue was sampled at anthesis and the third leaf was sampled at the third leaf stage (Zadoks 13-14), approximately 24 days before anthesis (Mutant, N = 8; Wild-type, N = 10). In 2016, one leaf was sampled per individual plant and was treated as an independent biological replicate. Similarly, in 2018, one leaf was sampled per row, and treated as an independent biological replicate. Three 1 cm^2^ discs were extracted from each leaf, one at the base of the leaf, one in the middle, and one from near the leaf tip. Chlorophyll was extracted as described previously (Wellburn, 1994); briefly, the discs of tissue were soaked in N,N-Dimethylformamide (analytical grade, Sigma Aldrich, UK) for 48-64 hrs until all pigment was completely removed from the leaf tissue. Pigment content was then quantified as previously described (Wellburn, 1994).

#### Leaf and Grain Mineral Content

Mineral content was taken from grain samples (2016) and leaf tissue samples (2018). Grain and leaf samples of approximately 0.2g were dried and ground to a fine powder before digestion with 2 mL nitric acid (67-69%, low-metal) and 0.5 mL hydrogen peroxide (30-32%, low-metal) for 12 hours at 95° C. Samples were then diluted 1:11 in ultrapure water before analysis with ICP-OES (Vista-PRO CCD Simultaneous ICP-OES; Agilent). Calibration was carried out using standards of Zn, Fe, and Mg at 0.2, 0.4, 0.6, 0.8, and 1 mg L^-1^ and Mn and P at 1,2, 3,4, and 5 mg L^-1^.

#### Light Microscopy

Thin sections of flag leaves were cut using a razor from mutant and wild-type plants in 2018 were imaged using a Leica MZ16 light microscope (Meyer Instruments, Houston, USA; N = 3 per genotype).

### Bulked Segregant Analysis

Individual plants with green and yellow phenotypes from the K2282 F_2_ populations sown at the JIC in 2016 were selected for bulked segregant analysis. DNA from plant tissues, sampled at seedling stage, was extracted using the QIAGEN DNeasy Plant Mini Kit. The quality and quantity of the DNA was checked using a DeNovix DS-11 Spectrophotometer, Qubit (High Sensitivity dsDNA assay, Q32854, Thermo Fisher), and by running a sample of the DNA on an agarose gel (1%) to visualise the high molecular weight DNA. Four bulks were assembled by pooling DNA from plants which had been scored as either “yellow” or “green” (K2282-28, N = 75 for yellow, N = 16 for green; K2282-23, N= 33 for yellow, N = 22 for green). Equal quantities of DNA from the individual plants were pooled into each bulk to minimise bias.

Library preparation and sequencing was carried out at the Earlham Institute (Norwich, UK) as follows. DNA quality control was carried out using the High Sensitivity Qubit assay, before library preparation was carried out with a KAPA HTP Library Prep Kit. Size selection was carried out using Beckman Coulter XP beads, and DNA was sheared to approximately 350 bp using the Covaris S2 sonicator. Four libraries were produced, one for each bulk detailed above, which were barcoded and pooled. Five cycles of PCR were carried out on the libraries before carrying out exome capture.

Hybridization to the wheat NimbleGen target capture, previously described in Krasileva *et al.* (2017), was carried out using the SeqCapEZ protocol v5.0, with the following changes: 2.8 μL of Universal Blocking Oligos was used, and the Cot-1 DNA was replaced with 14 μL of Developer Reagent. Hybridisation was carried out at 47°C for 72 hours in a PCR machine with a lid heated to 57°C.

The library pool was diluted to 2 nM with NaOH and 10μL transferred into 990μL HT1 (Illumina) to give a final concentration of 20pM. This was diluted further to an appropriate loading concentration in a volume of 120 μL and spiked with 1% PhiX Control v3 before loading onto the Illumina cBot. The flow cell was clustered using HiSeq PE Cluster Kit v4, utilising the Illumina PE_HiSeq_Cluster_Kit_V4_cBot_recipe_V9.0 method on the Illumina cBot. After clustering, the flow cell was loaded onto the Illumina HiSeq2500 instrument following the manufacturer’s instructions. The sequencing chemistry used was HiSeq SBS Kit v4. The library pool was run on two lanes with 125 bp paired end reads. Reads in bcl format were demultiplexed using the 6 bp Illumina index by CASAVA 1.8, allowing for a one base-pair mismatch per library, and converted to FASTQ format by bcl2fastq.

### Chromosome flow-sorting and sequencing

Seeds from the original K2282 M5 mutant line were used for the chromosome sorting and sequencing to ensure all parental SNPs were included. Suspensions of intact mitotic chromosomes were prepared from synchronized root tip meristems according to (Vrána et al., 2000). To achieve better discrimination of individual chromosomes by flow cytometry, GAA microsatellite loci were fluorescently labelled by FISHIS (Giorgi et al., 2013) using FITC-labelled (GAA)_7_ oligonucleotides as described (Vrána et al., 2016). Chromosomal DNA was then stained by 4’,6-diamidine-2’-phenylindole (DAPI) at final concentration 2 μg/ml and the chromosome suspensions were analysed by FACSAria SORP II flow sorter (BD Biosciences, San Jose, USA) at rates of 1000-2000 particles/sec. Bivariate flow karyotypes DAPI vs. GAA-FITC were obtained and individual populations were flow sorted to identify the population representing chromosome 3A and to estimate the extent of contamination by other chromosomes (Supplementary Figure 2). Briefly, 2000 chromosomes were sorted onto a microscopic slide and evaluated by fluorescence microscopy after FISH with probes for GAA microsatellite and Afa-family repeat (Kubaláková et al., 2002). Three batches of 30,000 copies of chromosome 3A corresponding to ~50 ng of DNA each were then sorted into PCR tubes containing 40 μl sterile deionized water. Chromosomal DNA was purified and amplified by Illustra GenomiPhi V2 DNA amplification Kit (GE Healthcare, Piscataway, USA) according to (Šimková et al., 2008).

Library preparation and sequencing were carried out at Novogene. DNA integrity was confirmed on 1% agarose gels. A PCR-free library preparation was carried out, using the NEBNext Ultra II DNA Library Prep Kit for Illumina, following manufacturer’s instructions. Libraries were sequenced using a HiseqX platform, generating 150 bp paired end reads.

### Sequencing Alignments and SNP calling

For the bulked segregant analysis, the raw Illumina reads were aligned to the Chinese Spring reference genome, RefSeqv1.0 (IWGSC et al., 2018), using bwa-mem (v 0.7.5) with the default settings (-k 20, -d 100) (Li, 2013). Alignments were sorted, indexed, and PCR duplicates removed using SAMtools (v 1.3.1) (Wysoker et al., 2009), and SNPs were called using freebayes (v 1.1.0, default settings) (Garrison and Marth, 2012). Depth of coverage was calculated using the exome capture size detailed previously (Krasileva et al., 2017) (Supplementary Table 2). Following SNP calling, we then filtered the original output to obtain only SNPs that were previously called in the K2282 parent line (Krasileva et al., 2017) using an original script available online to convert SNP coordinates to the RefSeq v1.0 genome (https://github.com/Uauy-Lab/K2282_scripts). The relative enrichment of each SNP in the yellow and green bulks was visualised across the wheat genome using the Circos package (Krzywinski et al., 2009). A schematic of the pipeline is provided in Supplemental Figure 3.

Following flow-sorting of chromosome 3A, reads were aligned to both RefSeq v1.0 and the Kronos assembly. We obtained access to the draft Kronos assembly produced at the Earlham Institute, which was assembled using the methods previously described (Clavijo et al., 2017a, Clavijo et al., 2017b). The Kronos assembly is available in advance of publication from Grassroots Genomics (https://opendata.earlham.ac.uk/opendata/data/Triticum_turgidum/). In both cases, the alignment was carried out with bwa-mem (v 0.7.5; default settings -k 20, -d 100) (Li, 2013). Illumina reads from the wild-type Kronos assembly were aligned to RefSeq v1.0 using hisat (v 2.0.4, default settings with -p 8) (Kim et al., 2015). In all cases, files were sorted, indexed, and PCR duplicates removed with SAMtools (v 1.3.1) (Wysoker et al., 2009). For alignments to RefSeq v1.0, depth of coverage across part 2 of chromosome 3A was calculated using genomic windows of 1 Mb (Supplementary Table 2). Depth of coverage was not calculated for the complete Kronos alignment, as the scaffolds are not associated with a chromosome. SNPs were called on the respective alignments using freebayes (v 1.1.0) at default settings in all cases. BCFtools (Wysoker et al., 2009) was used to filter the SNPs based on quality (QUAL ≥ 20), depth (DP > 10), zygosity (only homozygous), and EMS-like status (G/A or C/T SNPs). SNPs were also manually filtered to remove those which were likely to be varietal SNPs initially missed in filtering or which fell into regions of unexpectedly high SNP density. We then identified scaffolds from the Kronos genome which fall within the *YES-1* locus in the Chinese Spring RefSeq v1.0 genome using BLAST (v2.2.30) (Altschul et al., 1990) against the gene sequences annotated within that region, using the v1.1 gene annotation. All further analysis of the SNP data for mapping and marker design focused solely on the 32.9 Mb *YES-1* region. A schematic of this workflow is provided in Supplementary Figure 4.

### KASP Marker Genotyping

Markers were designed for the identified SNPs predominantly using the PolyMarker pipeline (Ramirez-Gonzalez et al., 2015b). Those not successful in PolyMarker were designed manually to be homoeolog specific. Markers were run on the recombinant populations using KASP genotyping, as previously described (Ramirez-Gonzalez et al., 2015a). Markers specific to K2282 are listed in Supplementary Table 3. Markers used for *NAM-A1* genotyping were previously published (Harrington et al., 2019).

### Data Analysis

Appropriate statistical tests for all data analyses were carried out and are detailed explicitly in the results section. When needed, adjustments for false discovery rate were carried out using the Benjamini-Hochberg adjustment. This is referred to in the results as “adjusted for FDR.” All statistics were carried out in R (v3.5.1) (R Core Team, 2018), and data was manipulated using packages tidyr (Wickham and Henry, 2018) and dplyr (Wickham et al., 2019). Graphs of phenotyping and expression data were produced using ggplot2 (Wickham, 2016) and gplots (Warnes et al., 2019), respectively.

## Results

### A forward screen of the Kronos TILLING population identifies a line segregating for a dominant chlorosis phenotype

951 M_4_ lines of the Kronos TILLING population (Krasileva et al., 2017) were grown at the John Innes Centre (JIC) in 2015 and scored for flag leaf and peduncle senescence timing. Ten lines showed early senescence phenotypes, while 11 showed late senescence phenotypes relative to Kronos wild-type (Supplementary Figure 5, Supplementary Table 1). We developed F_2_ populations for these 21 lines crossed to wild-type Kronos. In 2016 the F_2_ mapping populations for 18 of these 21 lines were grown at JIC, and again scored for the senescence. From these populations, two showed significantly delayed peduncle senescence; K1107, with delayed peduncle senescence present in two independent F_2_ populations, and K2711, with delayed peduncle senescence in one of two F_2_ populations (Supplementary Figure 6) These two lines both contained mutations in the *NAM-A1* gene, known to be a positive regulator of senescence (Uauy et al., 2006). The presence of the *NAM-A1* mutation was sufficient to account for the variation in peduncle senescence timing found within the F_2_ populations for both K1107 and K2711 (Tukey’s HSD, p < 0.01, Supplementary Figure 7), indicating that the *NAM-A1* SNPs were causal. The effect of the *NAM-A1* mutations was followed up separately (Harrington et al., 2019).

Based on the data from the 2016 field trials, we identified a single line, K2282, which showed a significant deviation in the timing of flag leaf senescence onset between the F_2_ population and the wild-type controls (p < 0.001, Kolmogorov-Smirnov test, adjusted for FDR; Supplementary Figure 6). Two F_2_ populations derived from K2282, K2282-28 and K2282-23, both showed earlier senescence compared to the wild-type controls. This phenotype, however, did not appear to be typical of a leaf senescence mutant. Although leaf senescence (scored based on leaf-tip necrosis) was indeed earlier in the K2282 populations, by anthesis the leaf tissue of individual plants was already highly chlorotic (Figure 1A). Quantification of chlorophyll levels confirmed that the yellow F_2_ individuals from both populations contained significantly less pigment than green F_2_ individuals (p < 0.05, Student’s t-test, Figure 1B). We also observed that the chlorosis phenotype predominated in the interveinal regions in the yellow plants, leading to a characteristic striated phenotype (Supplementary Figure 8).

We scored the K2282 F_2_ populations for chlorosis as a binary trait; *i.e.* plants were scored as yellow or green (see Fig. 1A for an image of yellow (MP/F_2_M) and green (WT) flag leaves). We confirmed that our visual scoring of the plants corresponded to the true chlorotic phenotype using non-destructive measurements of relative chlorophyll units. This identified a significant reduction in chlorophyll in the yellow (F_2_M) plants compared to the green (F_2_WT) plants, as expected (p < 0.001, Student’s t-test; Figure 1C). After classifying the F_2_ population into the green (F_2_WT) and yellow (F_2_M) groups, we found that the yellow group had significantly earlier leaf senescence (when scored to include necrotic symptoms) than the green group (p < 0.001 Kolmogorov-Smirnov test, Fig 1D). The segregation of the chlorotic phenotype within the two populations was not significantly different from a 3:1 yellow to green ratio (X^2^, p = 0.07; Figure 1E), consistent with the trait being underpinned by a single dominant locus, hereafter referred to as *Yellow Early Senescence 1 (YES-1)*.

### The *YES-1* locus maps to the long arm of chromosome 3A

To map the trait, we carried out bulked segregant analysis on the two independent populations, K2282-28 and K2282-23. A diagram of the analysis pipeline used is provided in Supplementary Figure 3. Following library preparation and exome capture, reads were aligned against the RefSeqv1.0 genome (IWGSC et al., 2018) and SNPs were called (Supplementary Table 2). To reduce the number of false SNP calls, we initially filtered the SNPs to only include those previously identified in the original M_2_ TILLING line (Krasileva et al., 2017). We recovered 1,548 SNPs out of the 3,060 SNPs present in the original K2282 M_2_ line which was sequenced. We expected to recover fewer SNPs than those identified in the original TILLING line as SNPs that were initially heterozygous in the M_2_ generation, may have been lost in the following two generations. Similarly, ~50% of heterozygous mutations present in the M_4_ line crossed to wild-type Kronos to produce the F_2_ population would also have been lost.

We initially focussed our analysis on the K2282-28 population and calculated the ratio of the mutant (alternate) allele over total depth of coverage (AO/DP) at each SNP location in the yellow and green bulks (Figure 2A, inner track). From this, we then calculated the Δ value representing the enrichment of the mutant allele in the yellow bulk compared to the green bulk (Figure 2A, outer track). The segregation ratio seen in the field suggested this is a dominant single locus trait. Hence, we assumed that the yellow bulk would contain individuals homozygous or heterozygous for the causal mutant allele, while the green bulk should only contain homozygous wild-type plants. As a result, the AO/DP value should approach 0 in the green bulk, and 0.66 in the mutant bulk, and thus have a Δ value of 0.66. Using a conservative limit of 0.5 for the Δ value (grey line, outer track of Figure 2A), we identified only one region, on Chromosome 3A, that was enriched for the mutant allele (Figure 2B). This result was consistent with that obtained from mapping carried out on the second population, K2282-23 (Supplementary Figure 9).

**Figure 2:**
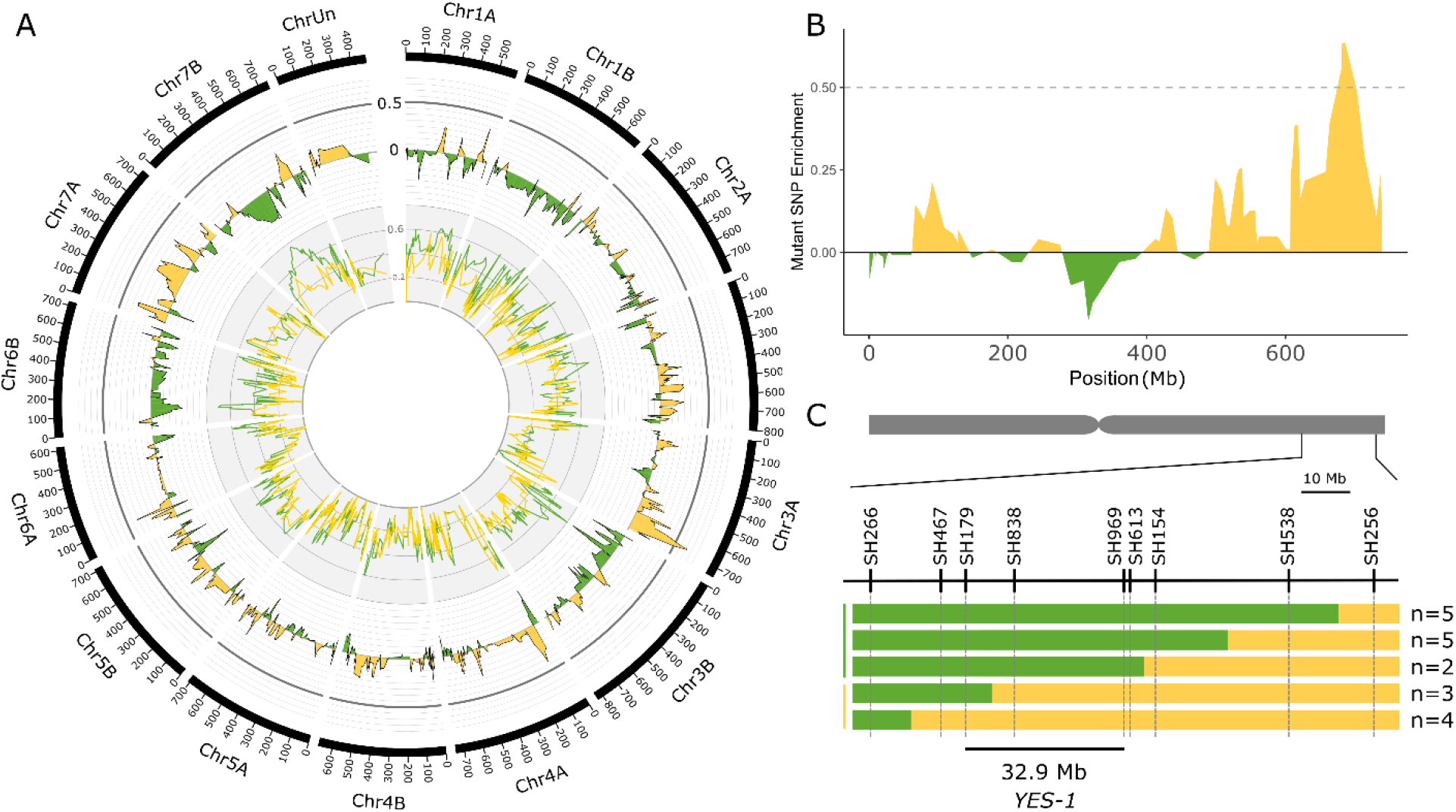
Bulked segregant analysis identifies the *YES-1* locus on chromosome 3A. Exome capture was carried out on yellow and green bulks from K2282 x Kronos F_2_ populations grown at the JIC in 2016. The K2282 yellow bulk (yellow line, inner track; smoothed to a moving average of 4) and green bulk (green line) were scored at each SNP locus identified for enrichment of the mutant allele. The level of enrichment in the green bulk was subtracted from that of the yellow bulk to obtain the Δ value (outer track; smoothed to a moving average of 4). A high Δ value, indicative of a region enriched for mutant alleles within the yellow bulks, was identified on the long arm of chromosome 3A (B; smoothed to a moving average of 4). Markers designed on known TILLING SNPs within this region mapped the *YES-1* locus to a 32.9 Mb interval within the F_2_ population (C). Green bars indicate wild-type calls, while yellow bars indicate mutant or heterozygous calls. The numbers of individual plants that fell into each recombination interval are shown to the right. The chromosome scale in (A) is given in Mb.

To validate this mapping, we developed KASP markers for the SNPs within and surrounding the region of interest (Figure 2C, Supplementary Table 3). Mapping of the individual F_2_ plants which were used to perform the exome capture confirmed the location of the region of interest on the long arm of chromosome 3A. Using the recombination events within this region and requiring at least two independent F_2_ plants to define the mapping interval, we narrowed the *YES-1* region to between markers SH179 and SH969, a region of 32.9 Mb in the RefSeq v1.0 genome containing 345 genes (RefSeq v1.1 gene annotation) (Figure 2C).

### Leaf chlorosis precedes anthesis but is inconsistent across environments

To further characterise the phenotype, individual lines which were genotyped as completely mutant or wild-type across the *YES-1* region were grown at the JIC in 2018. The mutant lines contained less chlorophyll A, B, and carotenoid pigment as early as the 3^rd^ leaf stage (Zadoks 13-14) (Student’s t-test, p < 0.01; Figure 3A). This difference was increased at anthesis (Student’s t-test, p < 0.005), at which stage there was a larger spread in pigment content within the mutant lines than the wild-type lines. Chlorophyll content, measured with SPAD units, was also monitored across the development of the plants, from 14 days before anthesis to 39 days post-anthesis. SPAD readings were consistently lower in the mutant lines up to 24 days post anthesis (p < 0.01, Pairwise Wilcoxon Rank Sum adjusted for FDR). The chlorotic phenotype remained highly visible on the leaves of the mutant plants, compared to wild-type (shown at 20 DPA, Figure 3C). In both wild-type and mutant lines, the level of chlorophyll in the flag leaf peaked at approximately 6 DPA (Figure 3B). No significant decline in SPAD units was observed in the wild-type plants until 24 DPA (p < 0.01, Pairwise Wilcoxon Rank Sum adjusted for FDR). In contrast, the mutant plants contained significantly less chlorophyll at 18 DPA compared to the peak at 6 DPA (p < 0.01, Pairwise Wilcoxon Rank Sum adjusted for FDR). Despite this earlier onset of senescence, the mutant lines continued to lose chlorophyll until the final stage of the time course (39 DPA), in line with the wild-type plants. We also found that the chlorosis phenotype is associated with significant decreases in leaf mineral content, with chlorotic leaves containing less magnesium at the 3^rd^ leaf stage, and less of all four measured minerals at anthesis (Mg, Fe, Zn, p < 0.05; Mn, p = 0.05; Supplementary Figure 10).

**Figure 3:**
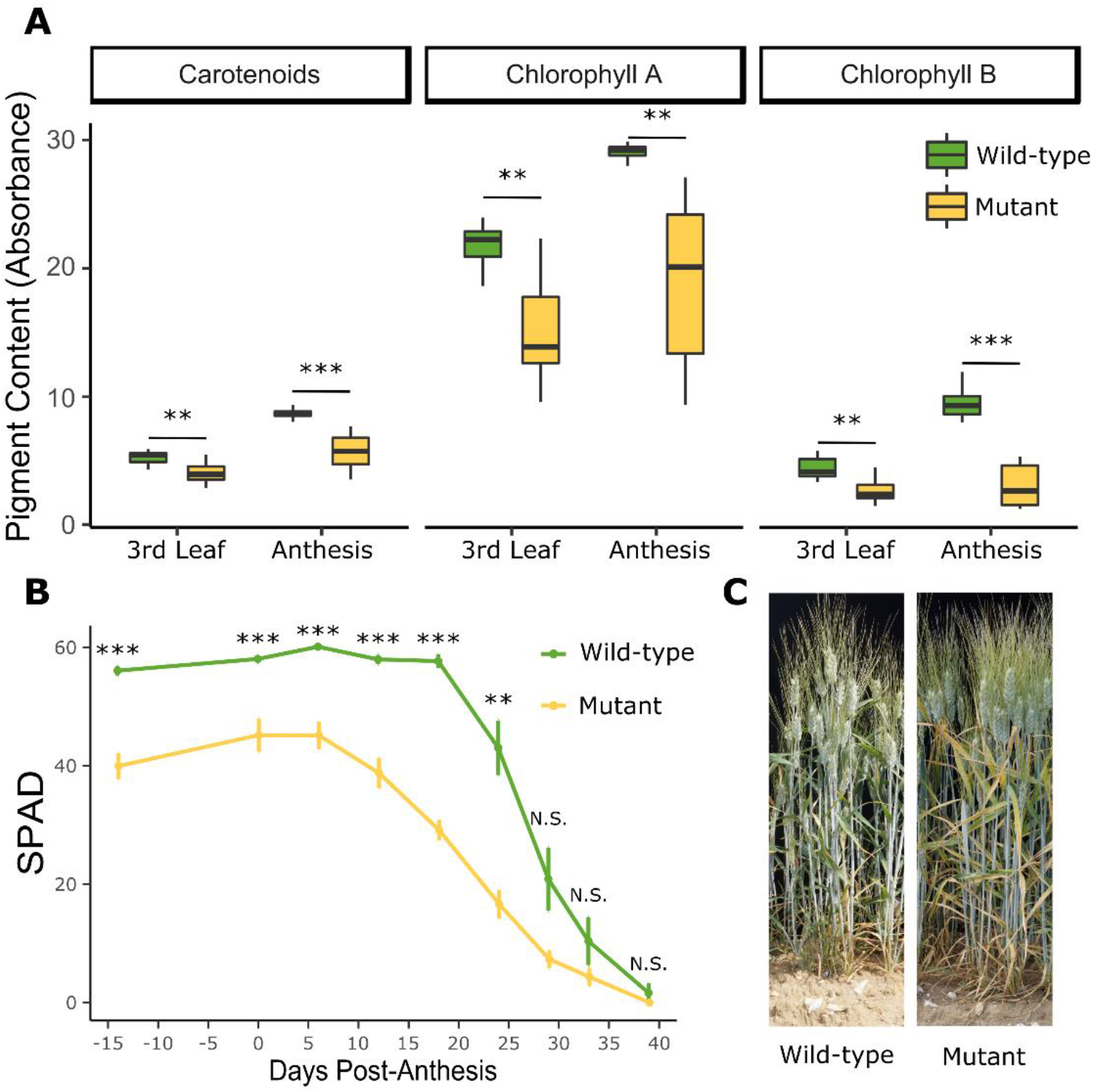
The *YES-1* locus causes lower chlorophyll levels before anthesis and earlier onset of senescence. The early chlorosis phenotype was recapitulated in the JIC 2018 field trials. Pigment content in the mutant lines is significantly lower at the third leaf stage (Zadoks 13-14, 24 days before anthesis) and becomes more extreme by anthesis (A; **, p < 0.01; ***, p < 0.001 Student’s T-test). Relatively chlorophyll content, as measured with a SPAD meter, is significantly decreased in the mutant lines before anthesis, and remains significantly lower until 29 days post-anthesis (B; **, p < 0.01; ***, p < 0.001 Pairwise Wilcoxon Rank Sum, adjusted for FDR). The yellowing phenotype in the leaves were clear in the field at 20 DPA (C).

Mutant and wild-type lines were also grown at UC Davis during the summer of 2016. Unlike in the UK, no chlorosis or senescence phenotype was observed either through visual or SPAD scoring (Supplementary Figure 11). This suggested that the causal locus underpinning *YES-1* is environmentally dependent. Given the similarity between the interveinal chlorosis phenotype observed in the *YES-1* mutant plants to that seen in plants with varying forms of nutrient deficiency (Snowball and Robson, 1991) and the decrease in leaf mineral content seen in the mutant plants (Supplementary Figure 8), we hypothesized that the environmental variation in phenotype may be due to nutrient content in the soil. To test this, F_3_ plants fully mutant across the *YES-1* region were grown under glasshouse conditions in three soil types: standard glasshouse cereal mix, soil taken from the JIC field site in 2017, and horticultural sand supplemented with nutrient-replete Hoagland solution. However, none of the three conditions tested recapitulated the yellowing phenotype observed in the field (Supplementary Figure 12). This was surprising given the consistency of the phenotype at the JIC field site across four different fields during four successive field seasons (2015-2018).

We then investigated weather-related environmental variation across the two field sites and across years. We obtained rainfall and temperature data from Davis, CA, for the 2016-2017 growing season, and from the JIC field site for the 2016, 2017, and 2018 growing seasons. The trials carried out in California in 2017 received substantially more rainfall between sowing and heading than in any of the JIC trials (Supplementary Table 4, Supplementary Figure 13). This suggested that perhaps reduced rain levels were correlated with the appearance of the mutant yellow phenotype. However, attempts to recapitulate the yellowing phenotype in the glasshouse through reduced watering of plants was also unsuccessful, as no early chlorosis or senescence was observed under different watering conditions (Supplementary Figure 12).

### Fine-mapping reduces the *YES-1* locus to 4 Mb on chromosome 3A

To identify further SNPs within the *YES-1* locus, we purified chromosome 3A from the K2282 mutant by flow cytometric sorting. However, as the population of 3A chromosomes partially overlapped with the population of 7A chromosomes on a bivariate flow karyotype DAPI vs. GAA-FITC (Supplementary Figure 2), flow-sorted fractions comprised 80% of chromosome 3A and 20% of chromosome 7A as determined by microscopic observation. For sequencing, three batches of 30,000 chromosomes (~50 ng) were flow-sorted and subsequent DNA amplification of three independent samples resulted in a total of 4.51 μg DNA.

Following sequencing, reads were mapped against the A and B genomes of the wheat RefSeq v1.0 genome. 60.38% of reads aligned to chromosome 3A while 25.37% aligned to chromosome 7A, consistent with the expected contamination. The remaining reads (14.25%) mapped against the rest of the genome. We obtained on average 82X coverage across chromosome 3A, using genomic windows of 1 Mb.

In order to maximise our ability to discover novel SNPs in the *YES-1* region, we carried out a simultaneous approach to SNP discovery utilising both the Chinese Spring reference genome as well as the draft Kronos assembly, as depicted in Supplementary Figure 4. In brief, paired end sequencing of the K2282 mutant chromosome 3A was used to obtain high-quality SNPs outside of the previously captured exome. We used the Kronos assembly to identify SNPs in non-coding regions that are less conserved between the Kronos and Chinese Spring cultivars. In tandem, we took advantage of the contiguity of the RefSeq v1.0 genome facilitated the identification of high-quality SNPs in and around all genes within the *YES-1* locus.

Reads from the mutant chromosome 3A were mapped against the draft Kronos assembly and were filtered for homozygous, EMS-like SNPs, passing minimum quality and depth thresholds. To obtain only SNPs that fell within the physical region encompassed by the *YES-1* locus, we carried out a BLAST between the Kronos scaffolds which contained SNPs and the Chinese Spring gene sequences within part of the *YES-1* region. Conducting a BLAST against gene sequences within the *YES-1* region, rather than the entirety of the region, reduced the number of scaffolds that mapped to the *YES-1* region due to shared repetitive sequences rather than true synteny. Based on recombination seen in individual plants, we focussed on a region encompassing markers SH179 and SH838, approximately 16 Mb in size. Within this region, we identified 18 unique Kronos scaffolds which both contained SNPs and at least one gene found in the RefSeqv1.0 *YES-1* physical interval (Supplementary Table 5). 26 of the genes within the *YES-1* region in Chinese Spring were identified (out of 345 total) within these 18 Kronos scaffolds. Genes that were not identified in the Kronos scaffolds may fall in scaffolds that contained no high-quality SNPs, may be split across multiple scaffolds, or may be absent from the Kronos genome. The SNPs within these scaffolds were manually curated, to exclude any regions that contained an unexpectedly high density of SNPs, leaving a final list of 16 scaffolds containing high-quality SNPs (Supplementary Table 5). The SNPs underlying markers SH838 and SH179, initially identified in the exome capture data, were also recovered in the Kronos genome, validating the use of this method. KASP primers were designed for a subset of the SNPs and were used, together with the previous phenotypic data, to map *YES-1* to a 6.6 Mb region between markers SH123480 and SH59985 (Figure 4A).

**Figure 4:**
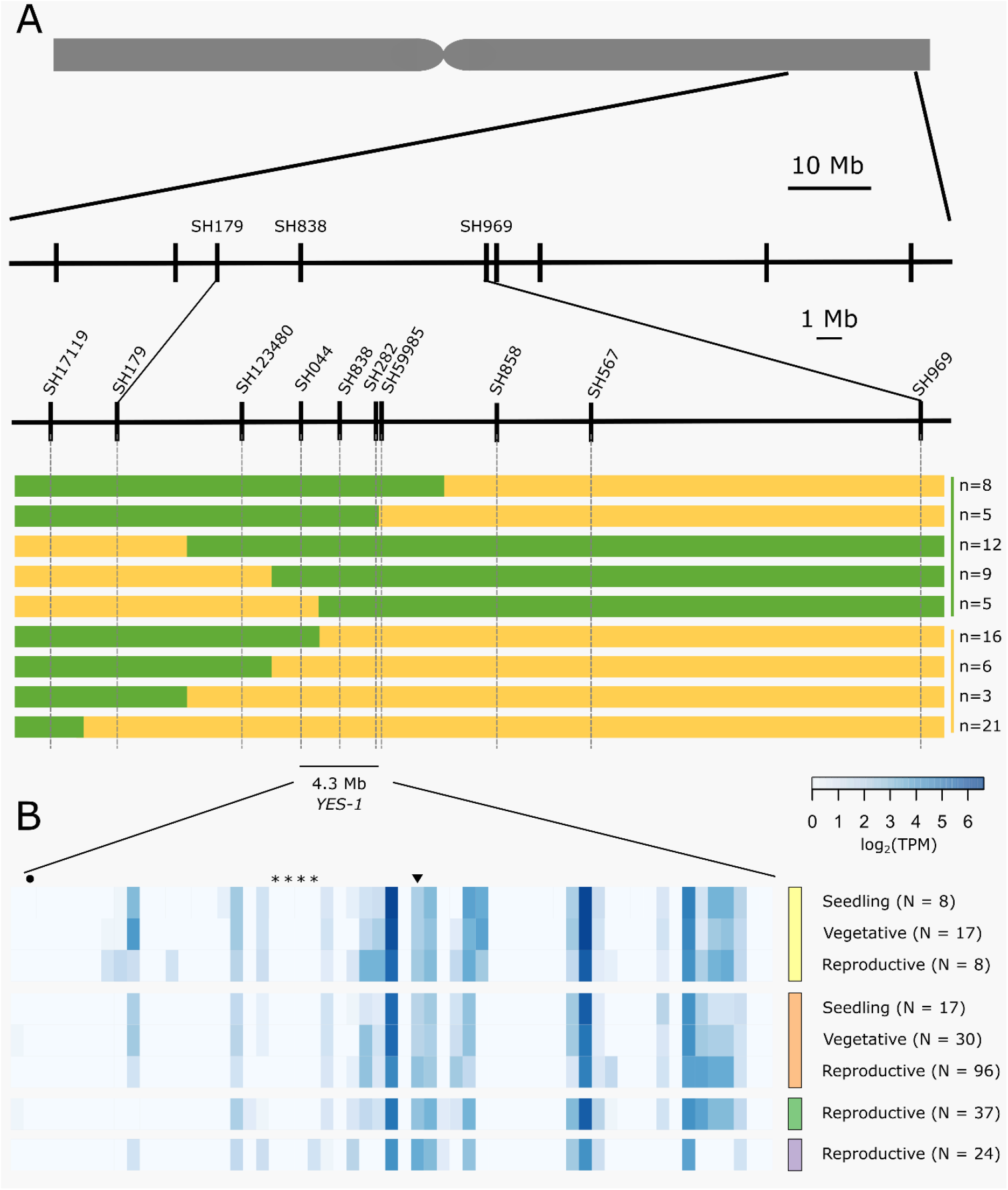
The *YES-1* locus fine-maps to a 4.3 Mb region containing 59 genes. Markers were developed for novel SNPs identified in non-coding regions. We used phenotypic data from JIC 2017 and 2018 field trials to classify recombinant lines as green or yellow. (A). These markers mapped *YES-1* to a 4.3 Mb interval between markers SH044 and SH59985. Expression data for the 59 high-confidence genes in the region (B) from developmental time course data (Ramírez-González et al., 2018) highlights gene expression in root (yellow, top), leaf/shoot (orange, second from top), spike (green, second from bottom) and grain (purple, bottom) tissues across developmental stages. Genes mentioned in the text are highlighted by an asterisk (TraesCS3A02G412900 to TraesCS3A02G413200; *OsSAG12* orthologs), a circle (TraesCS3A02G410800; *Tryptophan Decarboxylase 2)* or an inverted triangle (TraesCS3A02G414000; putative magnesium transporter).

To obtain more markers across the region, we also called SNPs against the Chinese Spring reference. To account for varietal SNPs between Kronos and Chinese Spring, we aligned raw reads from wild-type Kronos to the RefSeq v1.0 genome. Using a subset of reads, we obtained a coverage of approximately 30X across chromosome 3A. SNP calling was then carried out against RefSeq v1.0 to obtain a list of varietal SNPs between Chinese Spring and Kronos. A total of 968,482 homozygous SNPs with quality greater than 20 and depth of coverage greater than 10 were identified across the second part of chromosome 3A, encompassing *YES-1.*

SNPs were then called between the K2282 mutant chromosome 3A reads and the RefSeq v1.0 genome. The set of SNPs was filtered for quality and depth and to exclude the varietal SNPs identified above. Following this filtering, a total of 7,153 SNPs were identified between markers SH123480 and SH969, a region of approximately 30 Mb. This is substantially more SNPs than would be expected from the known mutation density of 23 mutations/Mb for the Kronos TILLING lines (Krasileva et al., 2017). However, SNP density across the region was highly irregular which we hypothesised was due to mismapping and spurious SNP calling in repetitive regions.

To reduce the impact of repetitive regions on SNP calling, we extracted SNPs only from regions encompassing 1 Kb up and downstream of annotated genes within the *YES-1* region. Following manual curation of SNPs, we identified a set of 15 SNPs that were located near genes within the annotated region (Supplementary Table 6). Of these SNPs, three were located in gene bodies (including the known TILLING SNP SH838), while the remainder were intronic (5) or fell in the promoter (5) or 3’ UTR (2). Note that some SNPs are in sufficient proximity to two gene models to be counted twice. Of the mutations in the coding region, SH838 and SH858 are both missense variants with low SIFT scores (0.00 and 0.03 respectively), while SH567 is a synonymous mutation. We designed markers based on these new SNPs and based on the JIC 2017 and 2018 phenotypic data we mapped *YES-1* to a 4.3 Mb interval, between markers SH044 and SH59985 (Figure 4A).

### Genes within the region

Within this region we identified 59 high-confidence genes based on the RefSeq v1.1 gene annotation (Supplementary Table 7). Using developmental time course data from two wheat varieties (Chinese Spring and Azhurnaya) (Borrill et al., 2016, Ramírez-González et al., 2018), we found that 25 genes within the region are expressed above 0.5 transcripts per million (TPM) in at least one stage of leaf or shoot tissue during development, consistent with our observation of a leaf-based phenotype (Figure 4). Of these genes, 18 were expressed above 0.5 TPM in leaf and shoot tissue during both vegetative and reproductive stages (Supplementary Table 7). This set of genes includes a putative magnesium transporter, TraesCS3A02G414000 (Gebert et al., 2009), which contains a missense mutation in the first exon of the gene which is predicted to be highly deleterious (SIFT = 0). This is the only gene within the 4 Mb region that contains a coding-region SNP, however no chlorosis phenotype was observed for any other line mutated in this gene (Supplemental Table 8). Within the 59 total candidate genes, five genes were found to have senescence-related functions in their closest rice orthologues. A set of four tandem duplicated genes, TraesCS3A02G412900 to TraesCS3A02G413200, are orthologues to the rice gene *OsSAG12-1,* a negative regulator of senescence (Singh et al., 2013). A fifth gene, TraesCS3A02G410800, is orthologous to *Tryptophan Decarboxylase 2*, a rice gene that causes higher serotonin levels and delayed leaf senescence when over-expressed (Kang et al., 2009). All five genes with senescence-related phenotypes are lowly expressed or non-expressed across the set of tissues and developmental stages considered. However, the majority of the genes within the region are un-annotated and lack orthologous copies in either rice or Arabidopsis.

## Discussion

Here we have fine-mapped a region causing a dominant, environmentally dependent early-chlorosis phenotype. We have taken advantage of the recently released genetic and genomic resources for wheat to increase our ability to identify SNPs *de novo* in a Kronos TILLING mutant line. We have shown how the use of cultivar specific genome assemblies can be used to increase the ability to identify high-quality SNPs in non-genic regions which are often relatively less conserved between varieties than coding sequences.

### Induced SNP variation can lead to novel dominant phenotypes

Many of the critical domestication alleles in polyploid wheat are derived from dominant mutations (Borrill et al., 2015, Uauy et al., 2017). This includes genes with critical variation in flowering time and free-threshing alleles resulting from dominant mutations (Fu et al., 2005, Yan et al., 2004, Simons et al., 2006, Greenwood et al., 2017). In wheat, the high level of redundancy between homoeologous genes adds to the importance of identifying dominant alleles to develop novel traits. Dominant alleles have retained their importance in modern breeding programs, underpinning the Green Revolution via the dominant dwarfing *Rht-1* allele (Peng et al., 1999, Borrill et al., 2015). Most traits selected for in modern breeding programs, however, lack standing variation of dominant alleles in both the modern breeding pool and in older wheat landraces and progenitors. Instead, forward screens for phenotypes of interest typically identify multiple quantitative trait loci (QTL) that each contribute towards a small portion of the desired phenotype. These more complex effects, often caused by loss of function mutations, are inherently more difficult to identify due to the need to acquire/combine mutations in both or all homoeologous copies of a gene to attain a clear phenotypic effect (Borrill et al., 2015, Borrill et al., 2019).

Here we have demonstrated that novel dominant alleles can be identified in chemically-mutagenised TILLING populations (Krasileva et al., 2017). Forward screens of the TILLING population are most likely to identify novel traits caused by dominant mutations, given the low likelihood of obtaining simultaneous mutations in multiple homoeologous copies of the same gene. Indeed, the fact that mutations in *NAM-A1* underpinned the only other senescence phenotype identified during this forward screen underscores this. The B-genome homoeolog of *NAM-A1* is non-functional in Kronos; as a result, a single mutation in the A-homoeolog equates to a complete null and was sufficient to show a strong and consistent phenotype (Pearce et al., 2014, Harrington et al., 2019).

The dominant phenotype identified in the K2282 line was particularly clear in that individual plants could unambiguously be scored for a binary green/yellow trait. However, we suggest that the TILLING population is equally well suited for forward genetic screens to identify novel dominant alleles governing other phenotypes. Recently, the Kronos TILLING population was used to identify a line which contained a deletion of *Rht-B1,* the partially-dominant dwarfing allele (Mo et al., 2018). Here we have identified a novel dominant allele with no previously characterised genes located within the candidate region. This highlights the potential for novel dominant alleles to be identified in populations with induced variation, such as the Kronos and Cadenza TILLING populations (Krasileva et al., 2017).

### The use of cultivar-specific assemblies facilitates the identification of non-genic SNPs

A complication of working with dominant induced variation, however, is that dominant mutations may often act through changes to regulatory elements. Variation in the promoter and intron sequence of the flowering time gene *VRN1* underpins the transition from winter to spring growth habit in wheat and barley (Yan et al., 2004, Fu et al., 2005). More recently, CRISPR editing has been used in tomato to edit the promoter region of various yield-related genes, leading to a high level of variation in trait morphology (Rodríguez-Leal et al., 2017). These results, amongst many others, highlight the potential importance of non-coding regions in regulating agronomically-relevant traits. However, many reduced representation methods focus on enrichment of coding regions (Borrill et al., 2019). Such methods of genome complexity reduction, therefore, are less likely to contain the information needed to identify a dominant causal SNP in a regulatory region. Compounding this difficulty is the fact that noncoding regions of the genome are typically less conserved between cultivars. As a result, SNP identification against the reference variety may fail to identify critical SNPs or, conversely, identify a large number of spurious SNPs.

We have shown here that the draft Kronos assembly can instead be used, alongside non-biased methods of genome size reduction (e.g. chromosome flow sorting), to identify cultivar-specific SNPs in non-coding regions of the genome. We started by calling SNPs against scaffolds of the Kronos assembly, obtaining a large amount of SNP variation between the wild-type and mutant lines. Once we had this data, we then positioned the scaffolds which contained SNPs against the reference genome, identifying the SNPs which were located within our region of interest (here *YES-1).* This approach overcame two of the main drawbacks to using the reference genome and the Kronos assembly. On one hand, the reference genome would be expected to have different sequence content to another variety, such as Kronos, limiting its utility for SNP identification. On the other hand, unlike the gold-standard reference genome, the Kronos assembly doesn’t have long-range assemblies needed to obtain positional information for SNPs. Long-range contiguous assemblies of additional cultivars, such as the recently published Svevo genome (Maccaferri et al., 2019), will greatly improve this current limitation. Until then, using variety-specific genomes, such as those being produced by the 10+ Wheat Genomes project, alongside the highly contiguous reference genomes will facilitate the identification of non-genic SNPs.

### Variability in phenotype points to an environmentally-dependent causal locus

The early chlorosis and senescence phenotype caused by the *YES-1* locus was consistent across four years in field trials at the JIC. However, mutant lines showed no evidence of a chlorotic phenotype when grown in Davis, CA. Comparison of rainfall and temperature patterns between the years and locations highlighted the fact that the plants received a high level of rainfall in Davis before flowering, substantially more than that received in any of the years at the JIC (Supplementary Table 4). This was due to the highly unusual wet winter that occurred in California in 2016/2017, with an average rainfall of 781 mm across the state from October 2016 to March 2017 (NOAA National Centers for Environmental Information, 2017). This suggested initially that the chlorosis response may be a response to higher water stress, yet we were unable to recapitulate the phenotype when grown in the glasshouse under different watering conditions.

We also considered whether the phenotype was due to variation in soil nutrient content. The presence of a missense mutation within the coding region of a putative Mg^2+^ transporter (Gebert et al., 2009) highlighted this as a promising candidate gene. Similarly, the observed interveinal chlorosis phenotype (Supplementary Figure 8) is reminiscent of that characteristic of a magnesium deficiency (Snowball and Robson, 1991). However, we failed to recapitulate the phenotype when grown in the glasshouse using soil taken from the field at JIC, and which should thus have the same nutrient composition. Compounding this, we found that Kronos TILLING lines which contained other SNPs within the transporter gene sequence did not show the same chlorotic phenotype (Supplementary Table 8). This included lines with both missense and premature stop codon mutations which lacked the exon containing the identified SNP in K2282. This implies that, if the magnesium transporter were the cause of the *YES-1* phenotype, the specific missense mutation present in K2282 has a unique ability to cause a dominant change in function. As the transporter is predicted to function in a hexamer, it is possible that the mutation could be sufficient to prevent the hexamer to function effectively once formed, but not sufficient to prevent the mutant monomer from being incorporated into the hexamer. In this way it may be possible that plants heterozygous for the mutation show an equally strong phenotype as homozygous mutants as incorporation of a mutant monomer disrupts completely the function of the hexameric complex. This hypothesis could be tested in the future using Cas9-driven base editing in wheat to recapitulate the exact mutation in an independent background (Zong et al., 2017).

An alternative possibility is that a separate SNP located in a regulatory region may be acting either on the identified magnesium transporter, or on a separate, currently uncharacterised gene. Few dominant chlorosis phenotypes have previously been reported in the literature. A dominant chlorosis phenotype was previously reported in *Brassica napus*, however this phenotype disappeared after budding unlike here, where the yellowing phenotype became increasingly strong post-heading (Wang et al., 2016). In wheat, a *Ygm* (yellow-green leaf colour) mutant has been identified with a semi-dominant phenotype where the heterozygous plants are an intermediate yellow-green colour between the wild-type and homozygous mutant plants (Wu et al., 2018). This phenotype is underpinned by abnormal chloroplast development and is associated with differential expression of genes involved in chlorophyll biosynthesis and carbon fixation, amongst other traits. Further work to fine-map the *YES-1* locus will hopefully shed light on the specific causal SNP underpinning the environmentally-dependent chlorosis phenotype observed here, as well as on mechanisms governing dominant traits in polyploid wheat.

## Acknowledgements

We would like to acknowledge A. Smith and M. Banfield (JIC) for helpful input during this project. We thank J. Vrána, M. Kubaláková and R. Šperková (IEB) for assistance with chromosome flow sorting and chromosome DNA amplification. The authors also thank J. Dubcovsky (UC Davis) for hosting the field trials at the University of California, Davis in 2016-2017. We also acknowledge the help of the field and glasshouse teams at the JIC and the University of California, Davis, as well as the help of G. Chilvers at the University of East Anglia ICP-OES platform. Finally, we thank B. Clavijo and the wheat pan-genome team at the Earlham Institute for allowing the use of the Kronos assembly before final publication.

## Author Contributions

CU and PB conceived the study; SAH, PB, and NC carried out the field trials and phenotyping; SAH carried out the mapping; MK and JD flow-sorted chromosome 3A determined the purity in flow-sorted fractions and amplified chromosomal DNA for sequencing. SAH and CU wrote the manuscript, and all authors have read and approved the final manuscript.

## Data Availability Statement

The raw reads from the exome capture and the flow-sorting experiments have been deposited on the SRA (PRJNA540141). The Kronos assembly is available from http://opendata.earlham.ac.uk/Triticum_turgidum/. Bespoke codes used for coordinate conversion are stored on Github (https://github.com/Uauy-Lab/K2282_scripts).

## References

10+ WHEAT GENOMES PROJECT. 2016. The Wheat ‘Pan Genome’ [Online]. Available: http://www.10wheatgenomes.com/ [Accessed 2019].

Acevedo-Garcia, J., Spencer, D., Thieron, H., Reinstädler, A., Hammond-Kosack, K., Phillips, A. L. & Panstruga, R. 2017. Mlo-based powdery mildew resistance in hexaploid bread wheat generated by a non-transgenic TILLING approach. Plant Biotechnology Journal, 15, 367–378.

Altschul, S. F., Gish, W., Miller, W., Myers, E. W. & Lipman, D. J. 1990. Basic local alignment search tool. Journal of Molecular Biology, 215, 403–410.

Avni, R., Zhao, R., Pearce, S., Jun, Y., Uauy, C., Tabbita, F., Fahima, T., Slade, A., Dubcovsky, J. & Distelfeld, A. 2014. Functional characterization of *GPC-1* genes in hexaploid wheat. Planta, 239, 313–324.

Borrill, P., Adamski, N. & Uauy, C. 2015. Genomics as the key to unlocking the polyploid potential of wheat. New Phytologist, 208, 1008–1022.

Borrill, P., Harrington, S. A., Simmonds, J. & Uauy, C. 2018. Identification of transcription factors regulating senescence in wheat through gene regulatory network modelling. bioRxiv, 456749.

Borrill, P., Harrington, S. A. & Uauy, C. 2019. Applying the latest advances in genomics and phenomics for trait discovery in polyploid wheat. The Plant Journal, 97, 56–72.

Borrill, P., Ramirez-Gonzalez, R. & Uauy, C. 2016. expVIP: a Customizable RNA-seq Data Analysis and Visualization Platform. Plant Physiology, 170, 2172.

Brinton, J. & Uauy, C. 2019. A reductionist approach to dissecting grain weight and yield in wheat. Journal of Integrative Plant Biology, 61, 337–358.

Clark, J. W. & Donoghue, P. C. J. 2018. Whole-Genome Duplication and Plant Macroevolution. Trends in Plant Science, 23, 933–945.

Clavijo, B. J., Garcia Accinelli, G., Wright, J., Heavens, D., Barr, K., Yanes, L. & Di-Palma, F. 2017a. W2RAP: a pipeline for high quality, robust assemblies of large complex genomes from short read data. bioRxiv, 110999.

Clavijo, B. J., Venturini, L., Schudoma, C., Accinelli, G. G., Kaithakottil, G., Wright, J., Borrill, P., Kettleborough, G., Heavens, D., Chapman, H., Lipscombe, J., Barker, T., Lu, F.-H., Mckenzie, N., Raats, D., Ramirez-Gonzalez, R. H., Coince, A., Peel, N., Percival-Alwyn, L., Duncan, O., Trösch, J., Yu, G., Bolser, D. M., Namaati, G., Kerhornou, A., Spannagl, M., Gundlach, H., Haberer, G., Davey, R. P., Fosker, C., Palma, F. D., Phillips, A. L., Millar, A. H., Kersey, P. J., Uauy, C., Krasileva, K. V., Swarbreck, D., Bevan, M. W. & Clark, M. D. 2017b. An improved assembly and annotation of the allohexaploid wheat genome identifies complete families of agronomic genes and provides genomic evidence for chromosomal translocations. Genome Research, 27, 885–896.

Dodsworth, S., Chase, M. W. & Leitch, A. R. 2016. Is post-polyploidization diploidization the key to the evolutionary success of angiosperms? Botanical Journal of the Linnean Society, 180, 15.

Doležel, J., Vrána, J., Šafář, J., Bartoš, J., Kubaláková, M. & Šimková, H. 2012. Chromosomes in the flow to simplify genome analysis. Functional & Integrative Genomics, 12, 397–416.

Dubcovsky, J. & Dvorak, J. 2007. Genome Plasticity a Key Factor in the Success of Polyploid Wheat Under Domestication. Science, 316, 1862.

Fu, D., Szűcs, P., Yan, L., Helguera, M., Skinner, J. S., Von Zitzewitz, J., Hayes, P. M. & Dubcovsky, J. 2005. Large deletions within the first intron in VRN-1 are associated with spring growth habit in barley and wheat. Molecular Genetics and Genomics, 273, 54–65.

Garrison, E. & Marth, G. 2012. Haplotype-Based Variant Detection From short-read sequencing. arXiv e-prints [Online]. Available: https://ui.adsabs.harvard.edu/abs/2012arXiv1207.3907G [Accessed July 01, 2012].

Gebert, M., Meschenmoser, K., Svidová, S., Weghuber, J., Schweyen, R., Eifler, K., Lenz, H., Weyand, K. & Knoop, V. 2009. A Root-Expressed Magnesium Transporter of the *MRS2/MGT* Gene Family in Arabidopsis thaliana Allows for Growth in Low-Mg^2+^ Environments. The Plant Cell, 21, 4018.

Giorgi, D., Farina, A., Grosso, V., Gennaro, A., Ceoloni, C. & Lucretti, S. 2013. FISHIS: Fluorescence In Situ Hybridization in Suspension and Chromosome Flow Sorting Made Easy. PLOS ONE, 8, e57994.

Greenwood, J. R., Finnegan, E. J., Watanabe, N., Trevaskis, B. & Swain, S. M. 2017. New alleles of the wheat domestication gene *Q* reveal multiple roles in growth and reproductive development. Development, 144, 1959.

Harrington, S. A., Overend, L. E., Cobo, N., Borrill, P. & Uauy, C. 2019. Conserved residues in the wheat (*Triticum aestivum*) NAM-A1 NAC domain are required for protein binding and when mutated lead to delayed peduncle and flag leaf senescence. bioRxiv, 573881.

Hoagland, D. R. & Arnon, D. I. 1950. The water-culture method for growing plants without soil. Circular. California Agricultural Experiment Station, 347, 32 pp.

Huang, X., Feng, Q., Qian, Q., Zhao, Q., Wang, L., Wang, A., Guan, J., Fan, D., Weng, Q., Huang, T., Dong, G., Sang, T. & Han, B. 2009. High-throughput genotyping by whole-genome resequencing. Genome Research, 19, 1068–1076.

IWGSC, Appels, R., Eversole, K., Stein, N., Feuillet, C., Keller, B., Rogers, J., Pozniak, C. J., Choulet, F., Distelfeld, A., Poland, J., Ronen, G., Sharpe, A. G., Barad, O., Baruch, K., Keeble-Gagnère, G., Mascher, M., Ben-Zvi, G., Josselin, A.-A., Himmelbach, A., Balfourier, F., Gutierrez-Gonzalez, J., Hayden, M., Koh, C., Muehlbauer, G., Pasam, R. K., Paux, E., Rigault, P., Tibbits, J., Tiwari, V., Spannagl, M., Lang, D., Gundlach, H., Haberer, G., Mayer, K. F. X., Ormanbekova, D., Prade, V., Šimková, H., Wicker, T., Swarbreck, D., Rimbert, H., Felder, M., Guilhot, N., Kaithakottil, G., Keilwagen, J., Leroy, P., Lux, T., Twardziok, S., Venturini, L., Juhász, A., Abrouk, M., Fischer, I., Uauy, C., Borrill, P., Ramirez-Gonzalez, R. H., Arnaud, D., Chalabi, S., Chalhoub, B., Cory, A., Datla, R., Davey, M. W., Jacobs, J., Robinson, S. J., Steuernagel, B., Van Ex, F., Wulff, B. B. H., Benhamed, M., Bendahmane, A., Concia, L., Latrasse, D., Bartoš, J., Bellec, A., Berges, H., Doležel, J., Frenkel, Z., Gill, B., Korol, A., Letellier, T., Olsen, O.-A., Singh, K., Valárik, M., Van Der Vossen, E., Vautrin, S., Weining, S., Fahima, T., Glikson, V., Raats, D., Číhalíková, J., Toegelová, H., Vrána, J., Sourdille, P., Darrier, B., Barabaschi, D., Cattivelli, L., Hernandez, P., Galvez, S., Budak, H., Jones, J. D. G., Witek, K., Yu, G., et al. 2018. Shifting the limits in wheat research and breeding using a fully annotated reference genome. Science, 361, eaar7191.

Jupe, F., Witek, K., Verweij, W., Sliwka, J., Pritchard, L., Etherington, G. J., Maclean, D., Cock, P. J., Leggett, R. M., Bryan, G. J., Cardle, L., Hein, I. & Jones, J. D. G. 2013. Resistance gene enrichment sequencing (RenSeq) enables reannotation of the NB-LRR gene family from sequenced plant genomes and rapid mapping of resistance loci in segregating populations. The Plant journal: for cell and molecular biology, 76, 530–544.

Kang, K., Kim, Y.-S., Park, S. & Back, K. 2009. Senescence-Induced Serotonin Biosynthesis and Its Role in Delaying Senescence in Rice Leaves. Plant Physiology, 150, 1380.

Kim, D., Langmead, B. & Salzberg, S. L. 2015. HISAT: a fast spliced aligner with low memory requirements. Nature Methods, 12, 357.

Krasileva, K. V., Vasquez-Gross, H. A., Howell, T., Bailey, P., Paraiso, F., Clissold, L., Simmonds, J., Ramirez-Gonzalez, R. H., Wang, X., Borrill, P., Fosker, C., Ayling, S., Phillips, A. L., Uauy, C. & Dubcovsky, J. 2017. Uncovering hidden variation in polyploid wheat. Proceedings of the National Academy of Sciences, 114, E913.

Krzywinski, M., Schein, J., Birol, İ., Connors, J., Gascoyne, R., Horsman, D., Jones, S. J. & Marra, M. A. 2009. Circos: An information aesthetic for comparative genomics. Genome Research, 19, 1639–1645.

Kubaláková, M., Vrána, J., Číhalíková, J., ŠImková, H. & DoležEl, J. 2002. Flow karyotyping and chromosome sorting in bread wheat (*Triticum aestivum L*.). Theoretical and Applied Genetics, 104, 1362–1372.

Li, H. 2013. Aligning sequence reads, clone sequences and assembly contigs with BWA-MEM. arXiv e-prints [Online]. Available: https://ui.adsabs.harvard.edu/abs/2013arXiv1303.3997L [Accessed March 01, 2013].

Maccaferri, M., Harris, N. S., Twardziok, S. O., Pasam, R. K., Gundlach, H., Spannagl, M., Ormanbekova, D., Lux, T., Prade, V. M., Milner, S. G., Himmelbach, A., Mascher, M., Bagnaresi, P., Faccioli, P., Cozzi, P., Lauria, M., Lazzari, B., Stella, A., Manconi, A., Gnocchi, M., Moscatelli, M., Avni, R., Deek, J., Biyiklioglu, S., Frascaroli, E., Corneti, S., Salvi, S., Sonnante, G., Desiderio, F., Marè, C., Crosatti, C., Mica, E., ÖZkan, H., Kilian, B., De Vita, P., Marone, D., Joukhadar, R., Mazzucotelli, E., Nigro, D., Gadaleta, A., Chao, S., Faris, J. D., Melo, A. T. O., Pumphrey, M., Pecchioni, N., Milanesi, L., Wiebe, K., Ens, J., Maclachlan, R. P., Clarke, J. M., Sharpe, A. G., Koh, C. S., Liang, K. Y. H., Taylor, G. J., Knox, R., Budak, H., Mastrangelo, A. M., Xu, S. S., Stein, N., Hale, I., Distelfeld, A., Hayden, M. J., Tuberosa, R., Walkowiak, S., Mayer, K. F. X., Ceriotti, A., Pozniak, C. J. & Cattivelli, L. 2019. Durum Wheat Genome Highlights Past Domestication Signatures And Future Improvement Targets. Nature Genetics.

Mamanova, L., Coffey, A. J., Scott, C. E., Kozarewa, I., Turner, E. H., Kumar, A., Howard, E., Shendure, J. & Turner, D. J. 2010. Target-enrichment strategies for next-generation sequencing. Nature Methods, 7, 111.

Mo, Y., Howell, T., Vasquez-Gross, H., De Haro, L. A., Dubcovsky, J. & Pearce, S. 2018. Mapping causal mutations by exome sequencing in a wheat TILLING population: a tall mutant case study. Molecular Genetics and Genomics, 293, 463–477.

NOAA NATIONAL CENTERS FOR ENVIRONMENTAL INFORMATION 2017. State of the Climate: National Climate Report for Annual 2017.

Paterson, A. H., Wang, X., Li, J. & Tang, H. 2012. Ancient and Recent Polyploidy in Monocots. In: P., S. & D., S. (eds.) Polyploidy and Genome Evolution. Berlin, Heidelberg: Springer.

Pearce, S., Tabbita, F., Cantu, D., Buffalo, V., Avni, R., Vazquez-Gross, H., Zhao, R., Conley, C. J., Distelfeld, A. & Dubcovksy, J. 2014. Regulation of Zn and Fe transporters by the GPC1gene during early wheat monocarpic senescence. BMC Plant Biology, 14, 368.

Peng, J., Richards, D. E., Hartley, N. M., Murphy, G. P., Devos, K. M., Flintham, J. E., Beales, J., Fish, L. J., Worland, A. J., Pelica, F., Sudhakar, D., Christou, P., Snape, J. W., Gale, M. D. & Harberd, N. P. 1999. ‘Green revolution’ genes encode mutant gibberellin response modulators. Nature, 400, 256–261.

R CORE TEAM 2018. R: A Language and Environment for Statistical Computing. In: Computing, R. F. F. S. (ed.). Vienna, Austria.

Ramürez-GonzáLez, R. H., Borrill, P., Lang, D., Harrington, S. A., Brinton, J., Venturini, L., Davey, M., Jacobs, J., Van Ex, F., Pasha, A., Khedikar, Y., Robinson, S. J., Cory, A. T., Florio, T., Concia, L., Juery, C., Schoonbeek, H., Steuernagel, B., Xiang, D., Ridout, C. J., Chalhoub, B., Mayer, K. F. X., Benhamed, M., Latrasse, D., Bendahmane, A., Wulff, B. B. H., Appels, R., Tiwari, V., Datla, R., Choulet, F., Pozniak, C. J., Provart, N. J., Sharpe, A. G., Paux, E., Spannagl, M., Bräutigam, A. & Uauy, C. 2018. The transcriptional landscape of polyploid wheat. Science, 361, eaar6089.

Ramirez-Gonzalez, R. H., Segovia, V., Bird, N., Fenwick, P., Holdgate, S., Berry, S., Jack, P., Caccamo, M. & Uauy, C. 2015a. RNA-Seq bulked segregant analysis enables the identification of high-resolution genetic markers for breeding in hexaploid wheat. Plant Biotechnology Journal, 13, 613–624.

Ramirez-Gonzalez, R. H., Uauy, C. & Caccamo, M. 2015b. PolyMarker: A fast polyploid primer design pipeline. Bioinformatics (Oxford, England), 31, 2038–2039.

Rodrüguez-Leal, D., Lemmon, Z. H., Man, J., Bartlett, M. E. & Lippman, Z. B. 2017. Engineering Quantitative Trait Variation for Crop Improvement by Genome Editing. Cell, 171, 470–480.e8.

Šimková, H., Svensson, J. T., Condamine, P., Hřibová, E., Suchánková, P., Bhat, P. R., Bartoš, J., Šafář, J., Close, T. J. & DoležEl, J. 2008. Coupling amplified DNA from flow-sorted chromosomes to high-density SNP mapping in barley. BMC Genomics, 9, 294.

Simons, K. J., Fellers, J. P., Trick, H. N., Zhang, Z., Tai, Y.-S., Gill, B. S. & Faris, J. D. 2006. Molecular Characterization of the Major Wheat Domestication Gene Q. Genetics, 172, 547.

Singh, S., Giri, M. K., Singh, P. K., Siddiqui, A. & Nandi, A. K. 2013. Down-regulation of *OsSAG12-1* results in enhanced senescence and pathogen-induced cell death in transgenic rice plants. Journal of Biosciences, 38, 583–592.

Snowball, K. & Robson, A. D. 1991. Nutrient Deficiencies and Toxicities in Wheat: A Guide for Field Identification. In: CIMMYT (ed.). Mexico, D.F.

Soltis, P. S. & Soltis, D. E. 2016. Ancient WGD events as drivers of key innovations in angiosperms. Current Opinion in Plant Biology, 30, 159–165.

Steuernagel, B., Periyannan, S. K., Hernández-Pinzón, I., Witek, K., Rouse, M. N., Yu, G., Hatta, A., Ayliffe, M., Bariana, H., Jones, J. D. G., Lagudah, E. S. & Wulff, B. B. H. 2016. Rapid cloning of disease-resistance genes in plants using mutagenesis and sequence capture. Nature Biotechnology, 34, 652.

Uauy, C. 2017. Wheat genomics comes of age. Current Opinion in Plant Biology, 36, 142–148.

Uauy, C., Distelfeld, A., Fahima, T., Blechl, A. & Dubcovsky, J. 2006. A NAC Gene Regulating Senescence Improves Grain Protein, Zinc, and Iron Content in Wheat. Science, 314, 1298.

Uauy, C., Wulff, B. B. H. & Dubcovsky, J. 2017. Combining Traditional Mutagenesis with New High-Throughput Sequencing and Genome Editing to Reveal Hidden Variation in Polyploid Wheat. Annual Review of Genetics, 51, 435–454.

Vrána, J., Cápal, P., ŠImková, H., Karafiátová, M., ČüžKová, J. & Doležel, J. 2016. Flow Analysis and Sorting of Plant Chromosomes. Current Protocols in Cytometry, 78, 5.3.1–5.3.43.

Vrána, J., Kubaláková, M., Simková, H., Číhalíkovái, J., Lysák, M. A. & Dolezel, J. 2000. Flow Sorting of Mitotic Chromosomes in Common Wheat (*Triticum aestivum* L.). Genetics, 156, 2033.

Vullo, A., Allot, A., Zadissia, A., Yates, A., Luciani, A., Moore, B., Bolt, B. J., Grabmueller, C., Ong, C. K., Bolser, D. M., Staines, D. M., Carvalho-Silva, D., Tapanari, E., Perry, E., Maslen, G., Williams, G., Naamati, G., Pedro, H., Sparrow, H., Allen, J. E., Howe, K. L., Taylor, K., Mcdowall, M. D., Russell, M., Barba, M., Paulini, M., Christensen, M., Kumar, N., Langridge, N., De Silva, N., Davis, P., Finn, R. D., Boddu, S., Potter, S., Maurel, T., Maheswari, U., Newman, V., Liu, Z., Kersey, P. J., Olson, A., Stein, J., Tello-Ruiz, M., Wei, S., Ware, D., Hammond-Kosack, K. E. & Urban, M. 2017. Ensembl Genomes 2018: an integrated omics infrastructure for non-vertebrate species. Nucleic Acids Research, 46, D802–D808.

Wang, W., Simmonds, J., Pan, Q., Davidson, D., He, F., Battal, A., Akhunova, A., Trick, H. N., Uauy, C. & Akhunov, E. 2018. Gene editing and mutagenesis reveal inter-cultivar differences and additivity in the contribution of *TaGW2* homoeologues to grain size and weight in wheat. Theoretical and Applied Genetics, 131, 2463–2475.

Wang, Y., He, Y., Yang, M., He, J., Xu, P., Shao, M., Chu, P. & Guan, R. 2016. Fine mapping of a dominant gene conferring chlorophyll-deficiency in *Brassica napus*. Scientific Reports, 6, 31419.

Warnes, G. R., Bolker, B., Bonebakker, L., Gentleman, R., Huber, W., Liaw, A., Lumley, T., Maechler, M., Magnusson, A., Moeller, S., Schwartz, M. & Venables, B. 2019. gplots: Various R Programming Tools for Plotting Data. R package version 3.0.1.1.

Wellburn, A. R. 1994. The Spectral Determination of Chlorophylls a and b, as well as Total Carotenoids, Using Various Solvents with Spectrophotometers of Different Resolution. Journal of Plant Physiology, 144, 307–313.

Wickham, H. 2016. ggplot2: Elegant Graphics for Data Analysis., New York, Springer-Verlag.

Wickham, H., François, R., Henry, L. & Müller, K. 2019. dplyr: A Grammar of Data Manipulation. R package version 0.8.0.1.

Wickham, H. & Henry, L. 2018. tidyr: Easily Tidy Data with ‘spread()’ and ‘gather()’ Functions. R package version 0.8.2.

Wu, H., Shi, N., An, X., Liu, C., Fu, H., Cao, L., Feng, Y., Sun, D. & Zhang, L. 2018. Candidate Genes for Yellow Leaf Color in Common Wheat (*Triticum aestivum* L.) and Major Related Metabolic Pathways according to Transcriptome Profiling. International Journal of Molecular Sciences, 19.

Wysoker, A., Handsaker, B., Marth, G., Abecasis, G., Li, H., Ruan, J., Homer, N., Durbin, R. & Fennell, T. 2009. The sequence Alignment/Map format and SAMtools. Bioinformatics, 25, 2078–2079.

Yan, L., Helguera, M., Kato, K., Fukuyama, S., Sherman, J. & Dubcovsky, J. 2004. Allelic variation at the *VRN-1* promoter region in polyploid wheat. Theoretical and Applied Genetics, 109, 1677–1686.

Yan, L., Loukoianov, A., Tranquilli, G., Helguera, M., Fahima, T. & Dubcovsky, J. 2003. Positional cloning of the wheat vernalization gene *VRN1*. Proceedings of the National Academy of Sciences, 100, 6263.

Zadoks, J. C., Chang, T. T. & Konzak, C. F. 1974. A decimal code for the growth stages of cereals. Weed Research, 14, 415–421.

Zong, Y., Wang, Y., Li, C., Zhang, R., Chen, K., Ran, Y., Qiu, J.-L., Wang, D. & Gao, C. 2017. Precise base editing in rice, wheat and maize with a Cas9-cytidine deaminase fusion. Nature Biotechnology, 35, 438.

