## Supplementary Files for "Identification of a dominant chlorosis phenotype through a forward screen of the *Triticum turgidum* cv. Kronos TILLING population"

**Supplementary File**

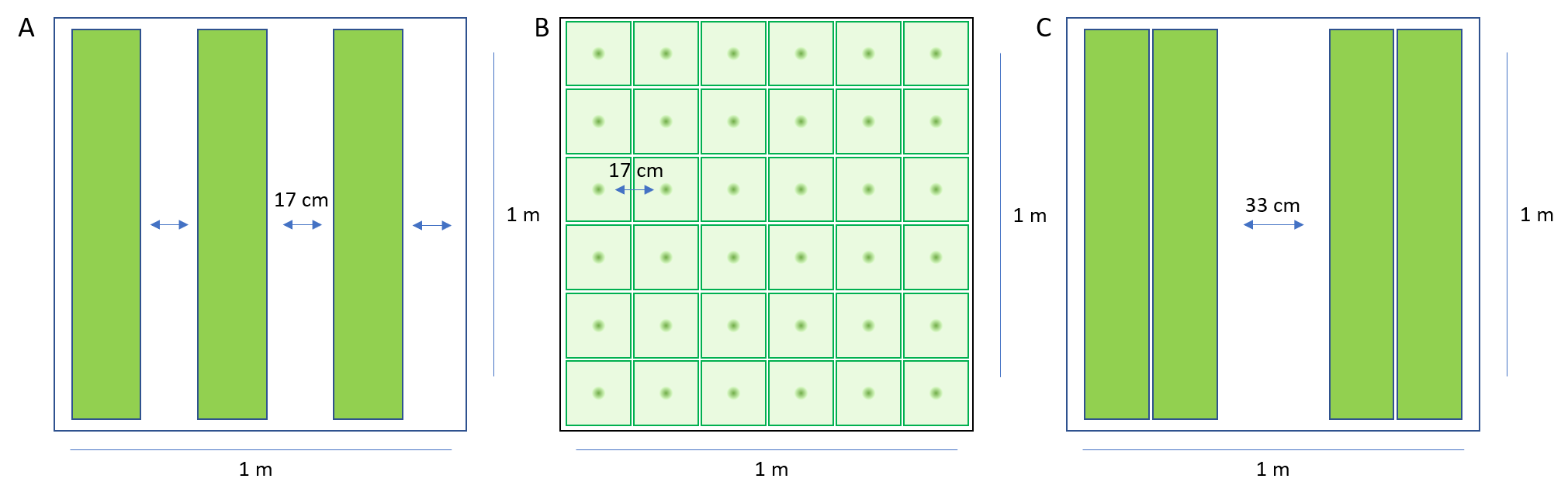

**Supplementary Figure 1: Diagram of field trial plot layouts.** The 2015 and 2017 JIC trials were laid out as in (A), with three single 1 m rows of plants (green) sown per 1 m^2^ plot, separated by approximately 17 cm. The 2016 JIC trials were hand-sown into a 6x6 grid in the 1 m^2^ plot (B), with approximately 17 cm between each individual plant. The 2018 JIC trial and the 2017 Davis, CA trial were both sown with 2 double rows per 1 m^2^ plot, each separated by approximately 33 cm (C).

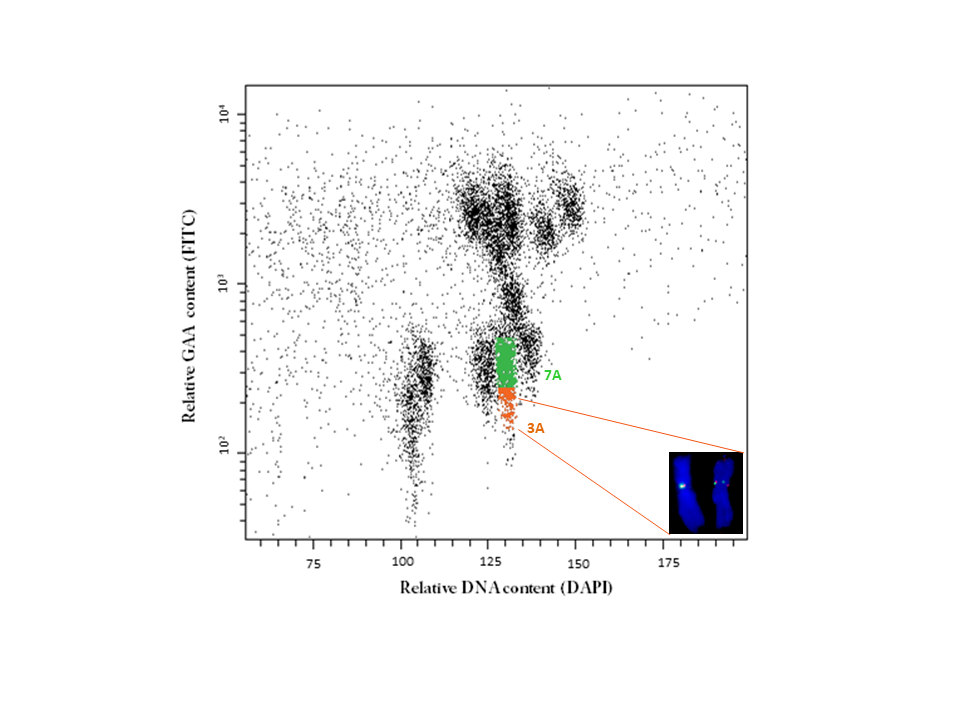

**Supplementary Figure 2: Bivariate flow karyotype DAPI vs. GAA-FITC obtained after the analysis of mitotic chromosomes isolated from *Triticum durum* cv. Kronos.** The regions representing chromosomes 3A and 7A are highlighted in orange and green, respectively. Inset: Images of flow-sorted chromosome 3A, which was identified by FISH with probes for GAA microsatellite (green) and Afa-family repeat (red). The chromosomes were counterstained by DAPI (blue).

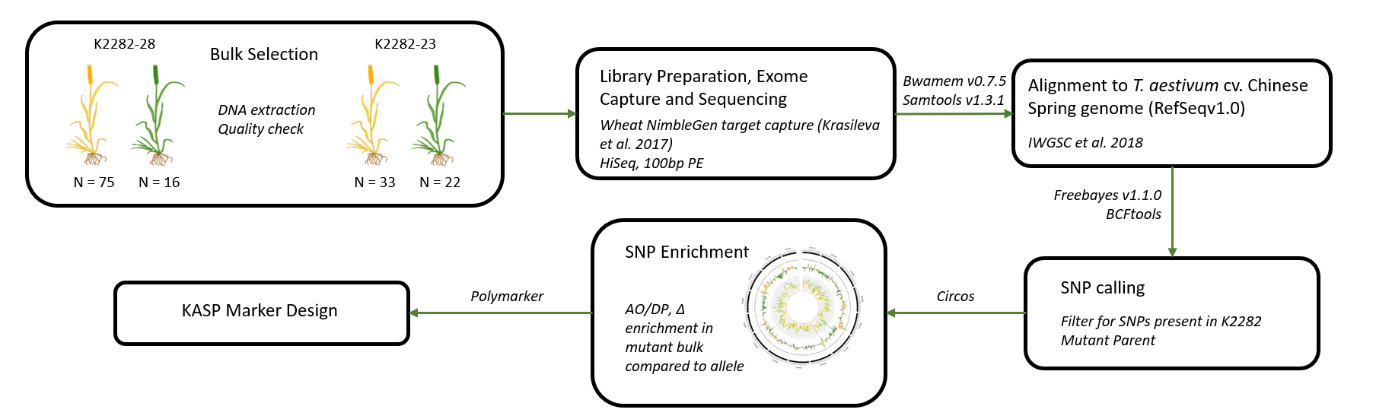

**Supplementary Figure 3: Pipeline for analysis of bulked segregant exome capture data.** Exome capture was carried out on pooled DNA from yellow and green plants in two independent F_2_ populations (K2282-23 and K2282-28). The enrichment of SNPs in the yellow bulks, which were derived from the original mutant parent line (K2282), was used to identify a region of interest on chromosome 3A. See methods for more details.

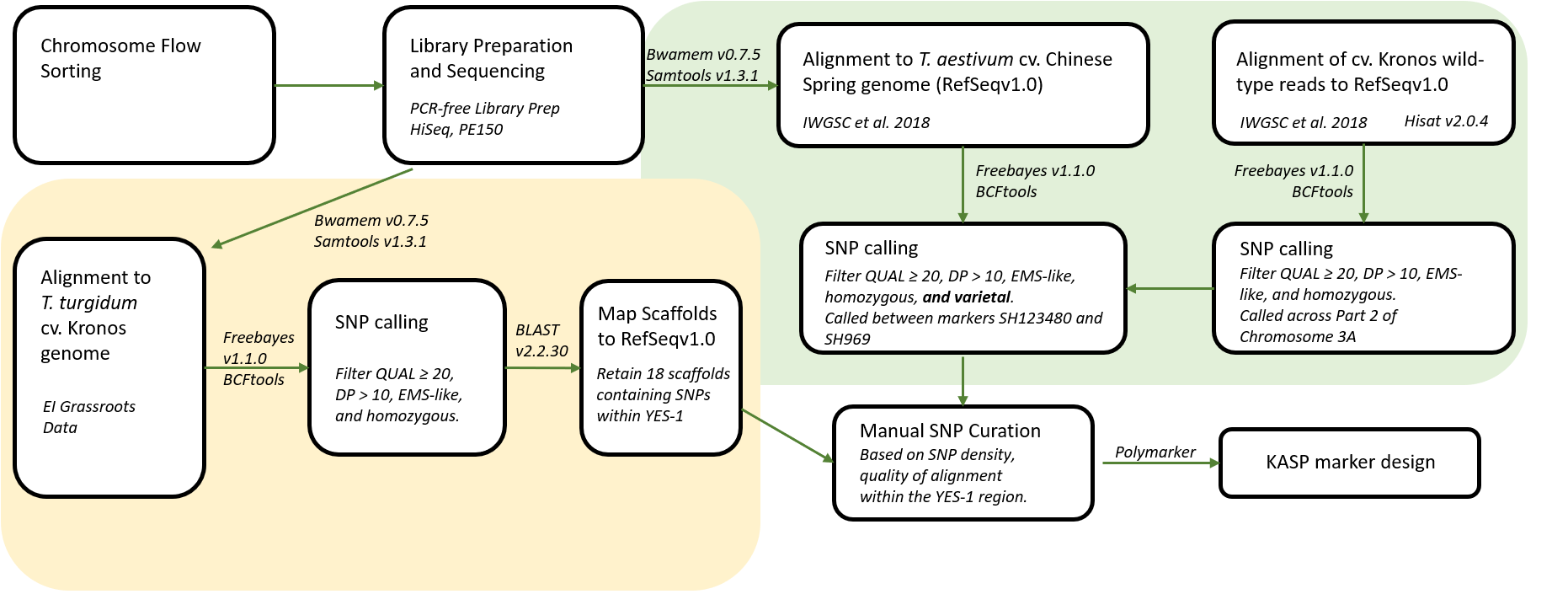

**Supplementary Figure 4: Pipeline for analysis of chromosome flow-sorting data.** DNA obtained from the purified Kronos2282 chromosome 3A was sequenced using Illumina sequencing. Reads were processed in two parallel paths; aligned to the cultivar Kronos assembly (yellow) and aligned to the reference Chinese Spring genome (green). SNPs obtained from both methods were used for KASP marker design. Bioinformatics tools used in each step are shown next to the corresponding arrow.

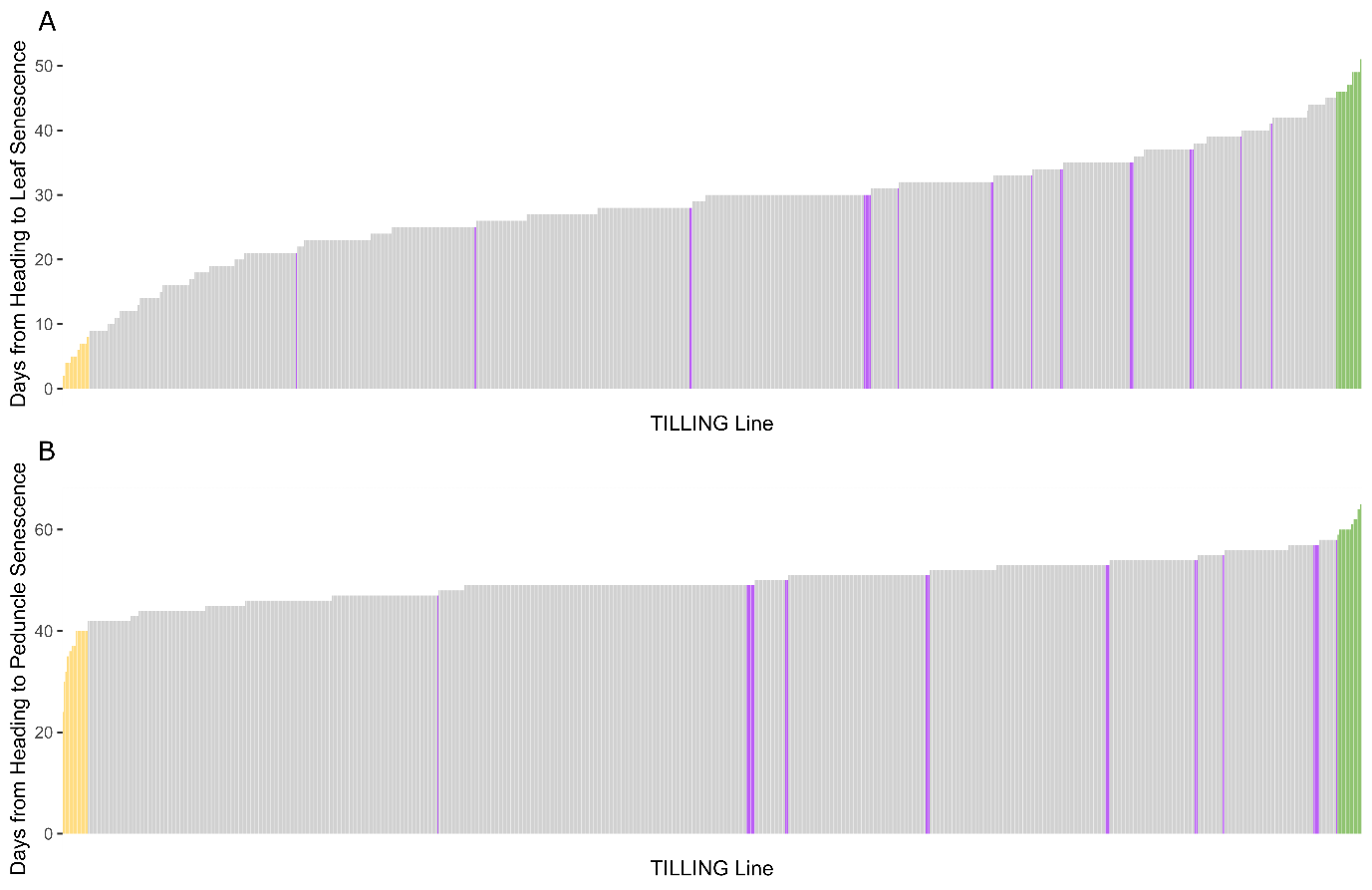

**Supplementary Figure 5: Early and late senescing TILLING lines were identified in a forward screen.** 951 TILLING lines from the Kronos TILLING population were screened for leaf (A) and peduncle (B) senescence onset. The 10 earliest senescing lines are highlighted in yellow, and the 11 latest senescing lines are highlighted in green. Wild-type Kronos was sown as a control, and the senescence timings for the wild-type replicates are shown in purple.

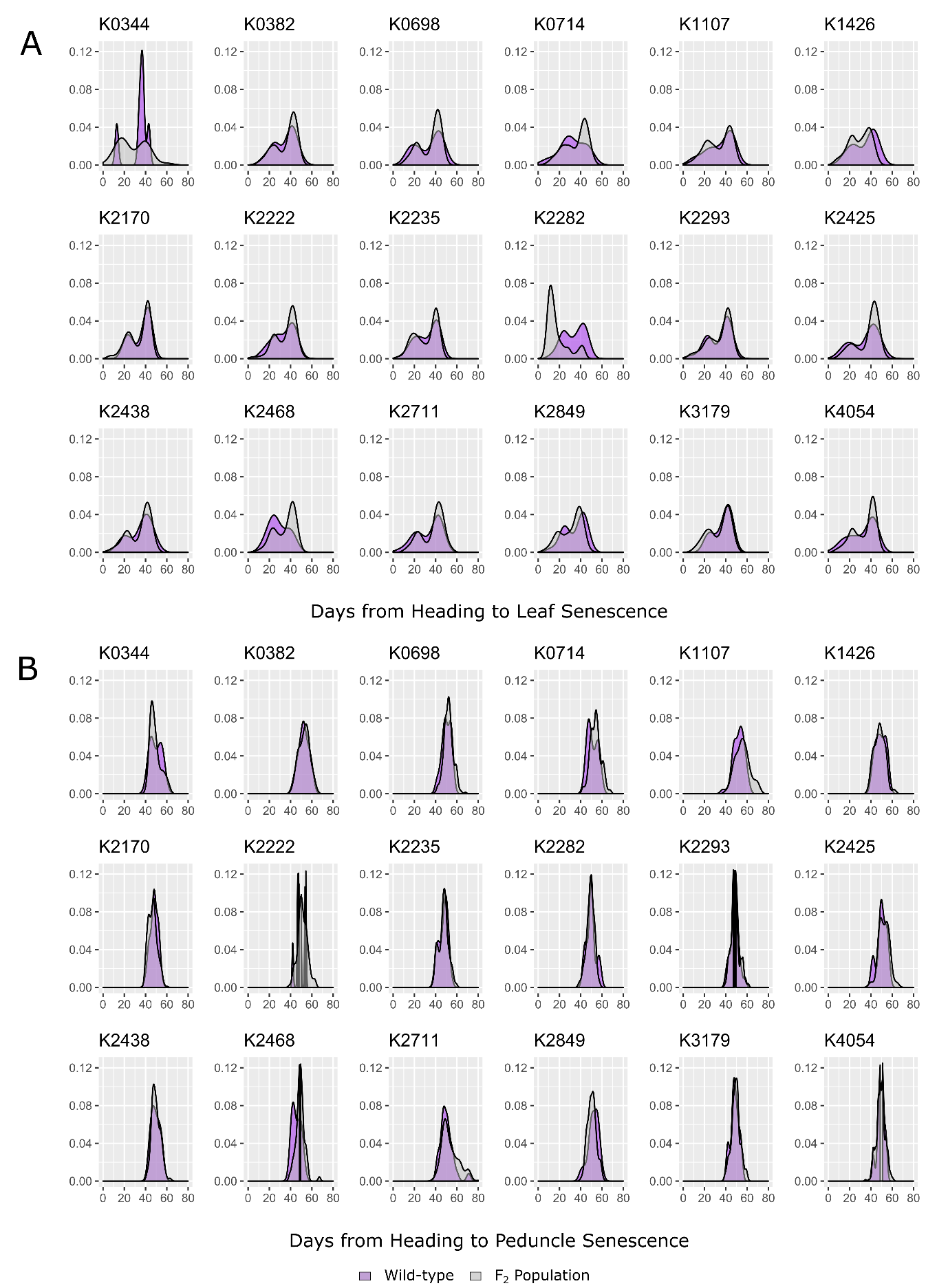

**Supplementary Figure 6: Segregating F_2­_ populations derived from Kronos TILLING lines show early and late senescence phenotypes.** Leaf senescence (A) and peduncle senescence (B) was scored for individual plants in F_2_ populations at the JIC in 2016. The distribution of senescence timings for Kronos wild-type control (purple) and the F_2_ individuals (grey) are shown. The F_2_ population only differs significantly from the wild-type for leaf senescence in line K2282 (p < 0.001, Kolmogorov Smirnov test adjusted for FDR). Note that the wild-type distributions derive from wild-type plants grown within the respective population; as a result, the wild-type distributions will differ between the different populations.

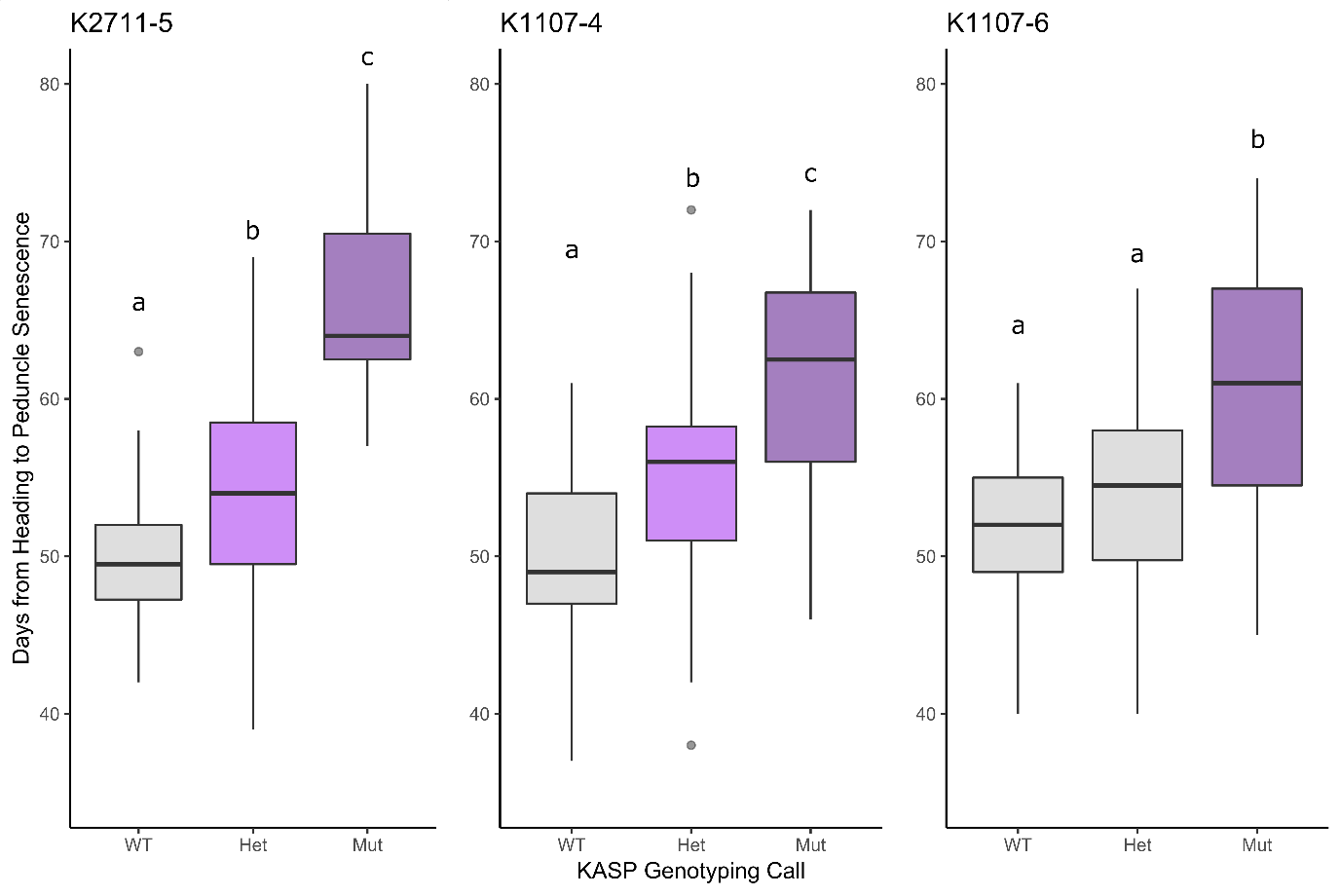
**Supplementary Figure 7: Mutations in *NAM-A1* explain the delay in peduncle senescence in K2711 and K1107.** Markers specific to the *NAM-A1* mutations in K1107 and K2711 were used to genotype the F_2_ populations in 2016. Significant delays in senescence (p < 0.01, Tukey’s HSD) are indicated with letters; boxplots sharing the same letter (as in K1107-6) are not significantly different in senescence timing. Note that the second K2711 population, K2711-6, was not significantly delayed in senescence and did not contain the *NAM-A1* mutation (data not shown). N ranged from 27 (K2711 Mut) to 116 (K1107-6 Het).

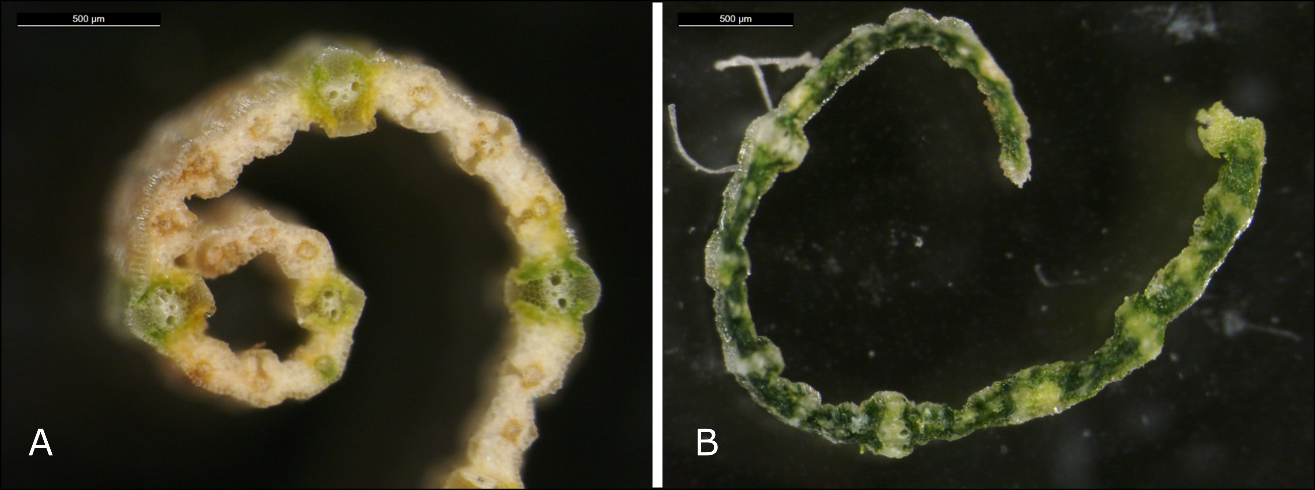

**Supplementary Figure 8: K2282 mutants show a striated inter-veinal chlorosis phenotype.** Mutant plants (A) showed a characteristic loss of pigment in the inter-veinal regions, retaining green colouration in the main veins of the flag leaf, compared to the wild-type plants (B).

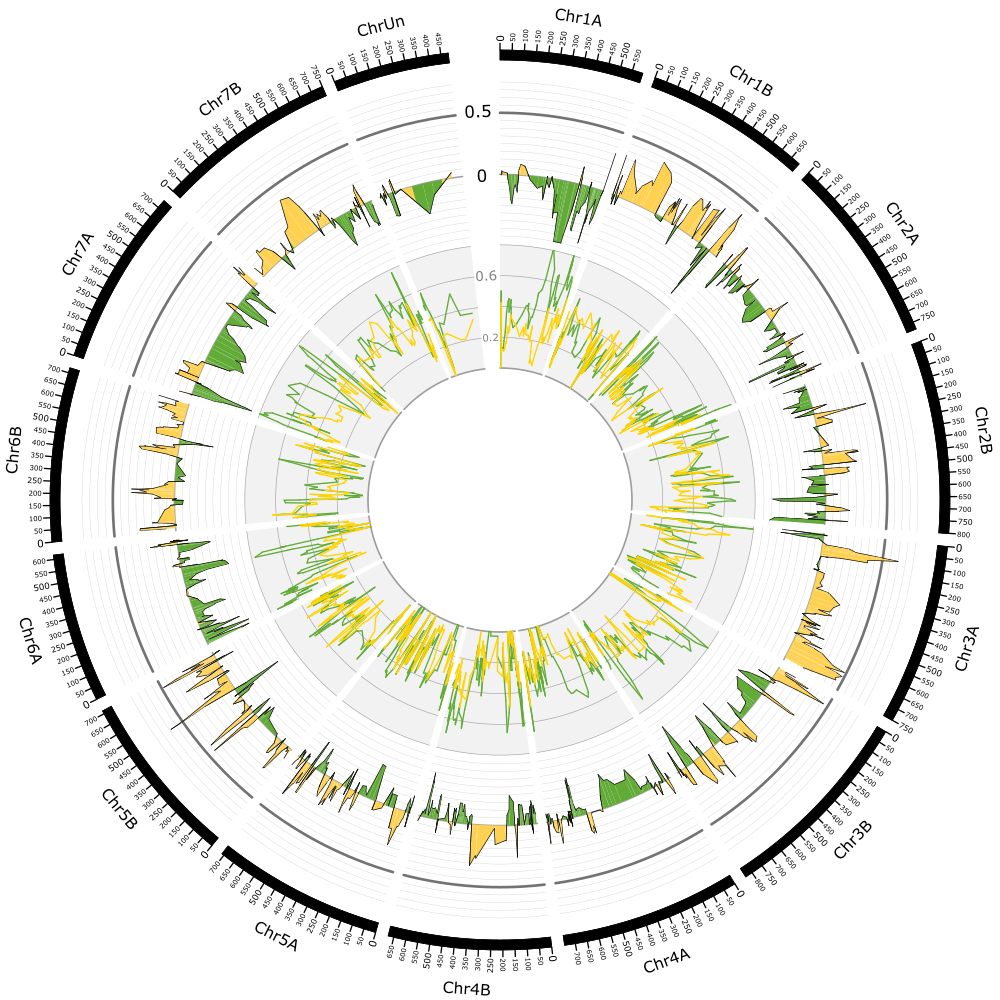

**Supplementary Figure 9: Genome-wide enrichment for mutant SNPs from the K2282-23 population.** The enrichment of the mutant allele at each SNP position (AO/DP) is shown on the inner track, with the green bulk highlighted in green and the yellow bulk highlighted in yellow. The scale for the inner track is shown between ChrUn and Chr1A in grey. The Δ value at each SNP position is shown in the outer track, with regions enriched for the mutant allele shown in yellow, and those enriched for the wild-type allele shown in green. The scale for the outer track is shown in black between ChrUn and Chr1A. The cut-off of 0.5 for the Δ value is shown as a thick grey line in the outer track. In both tracks, the trend shown is the moving average with an interval of 4. The Δ value approached the 0.5 cut-off in both the K2282-23 and K2282-28 (Figure 2 main text) populations only at the end of chromosome 3A. The chromosome scale is presented in Mb.

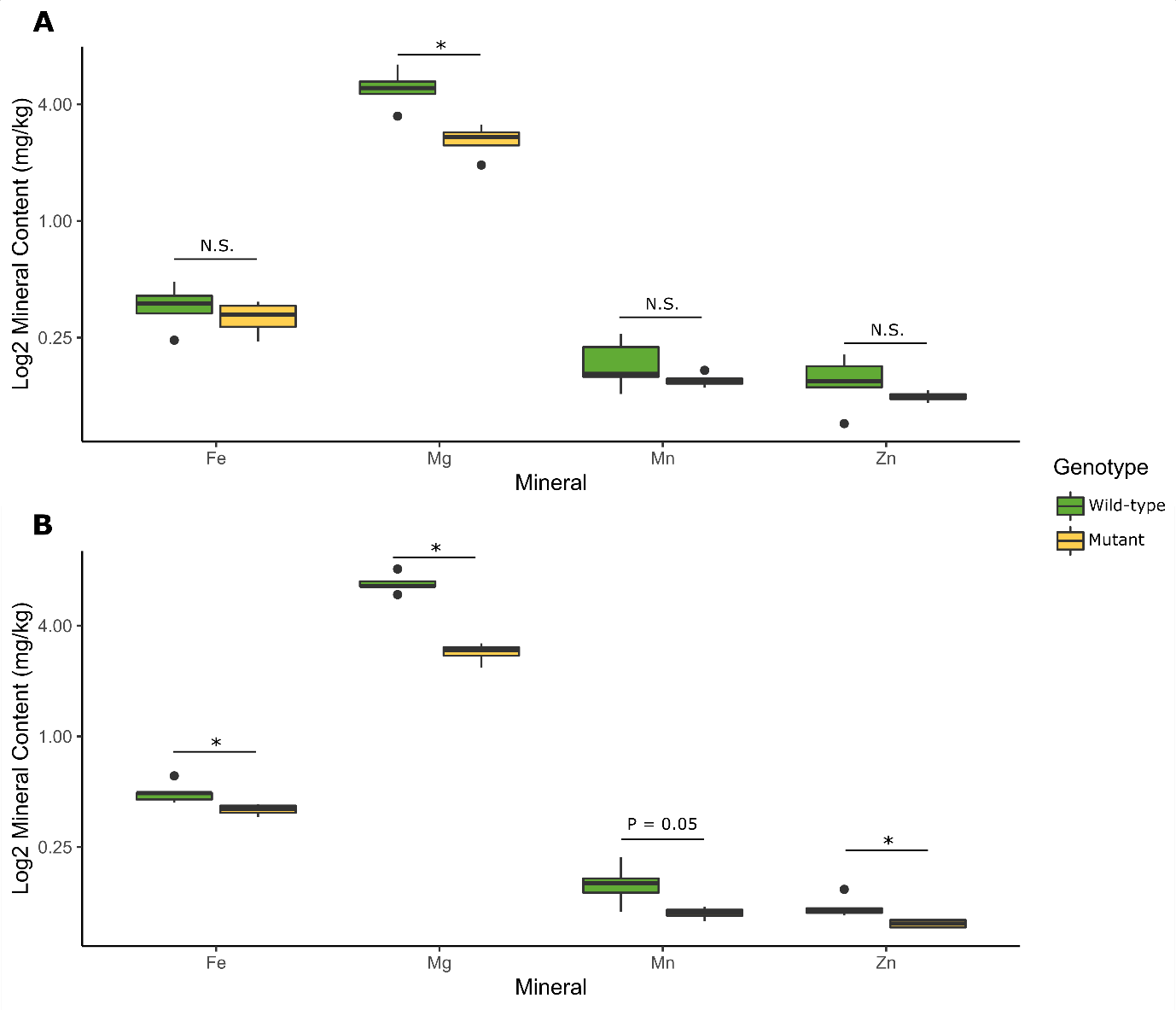

**Supplementary Figure 10: Mutant plants have lower mineral content.** Measurements of leaf mineral content were carried out at the 3^rd^ leaf stage (A; Zadoks 13-14) and anthesis (B) for wild-type and mutant plants at the JIC in 2018.

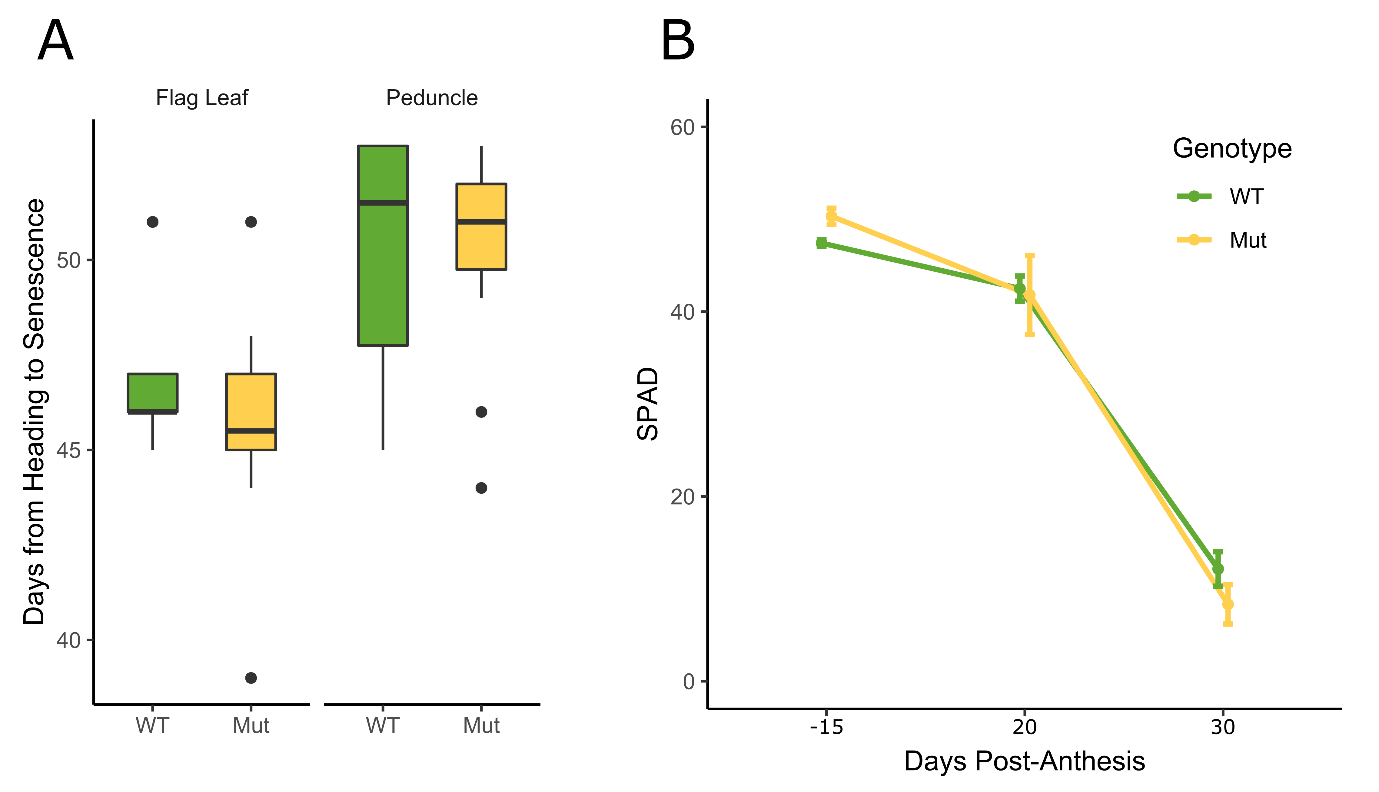

**Supplementary Figure 11: No visual senescence of chlorosis was observed in Davis, CA**. Rows of F_3_ individuals descended from individual F_2_ plants which were either fully wild-type or mutant across the *YES-1* region were phenotyped for visual senescence (A) and relative chlorophyll units using SPAD (B). No significant difference in senescence or chlorosis was observed at any stage.

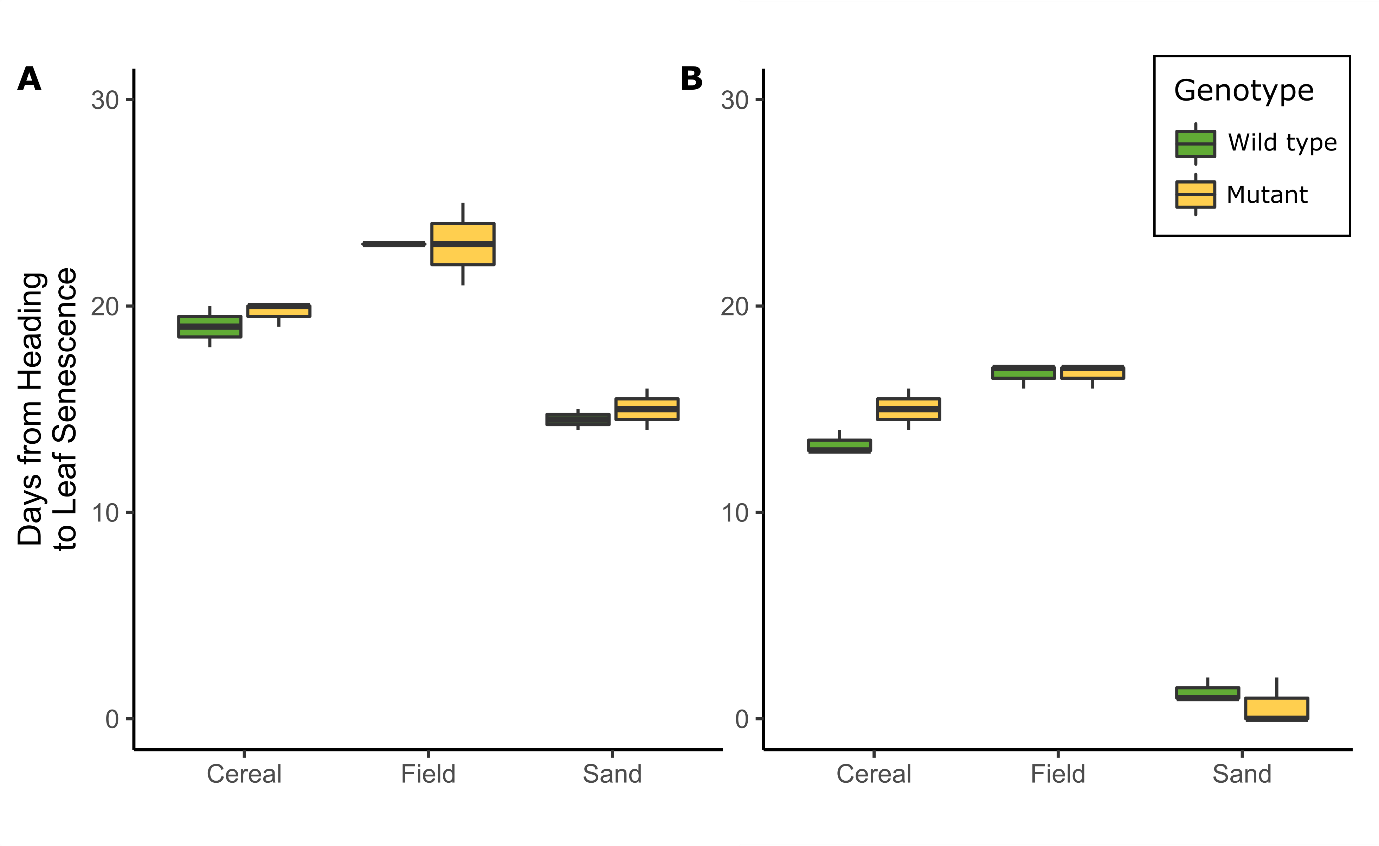

**Supplementary Figure 12: Soil and drought conditions could not recapitulate the early chlorosis phenotype in the glasshouse.** Mutant and wild-type plants were grown under glasshouse conditions in three soil conditions (Cereal Mix, JIC 2017 Field soil, and Horticultural Sand supplemented with Hoagland Solution) and under normal (A) and low water (B) conditions. No significant early onset of chlorosis or senescence was observed for the mutant plants in any condition (Student’s t-test). N=3 for each genotype and condition combination.

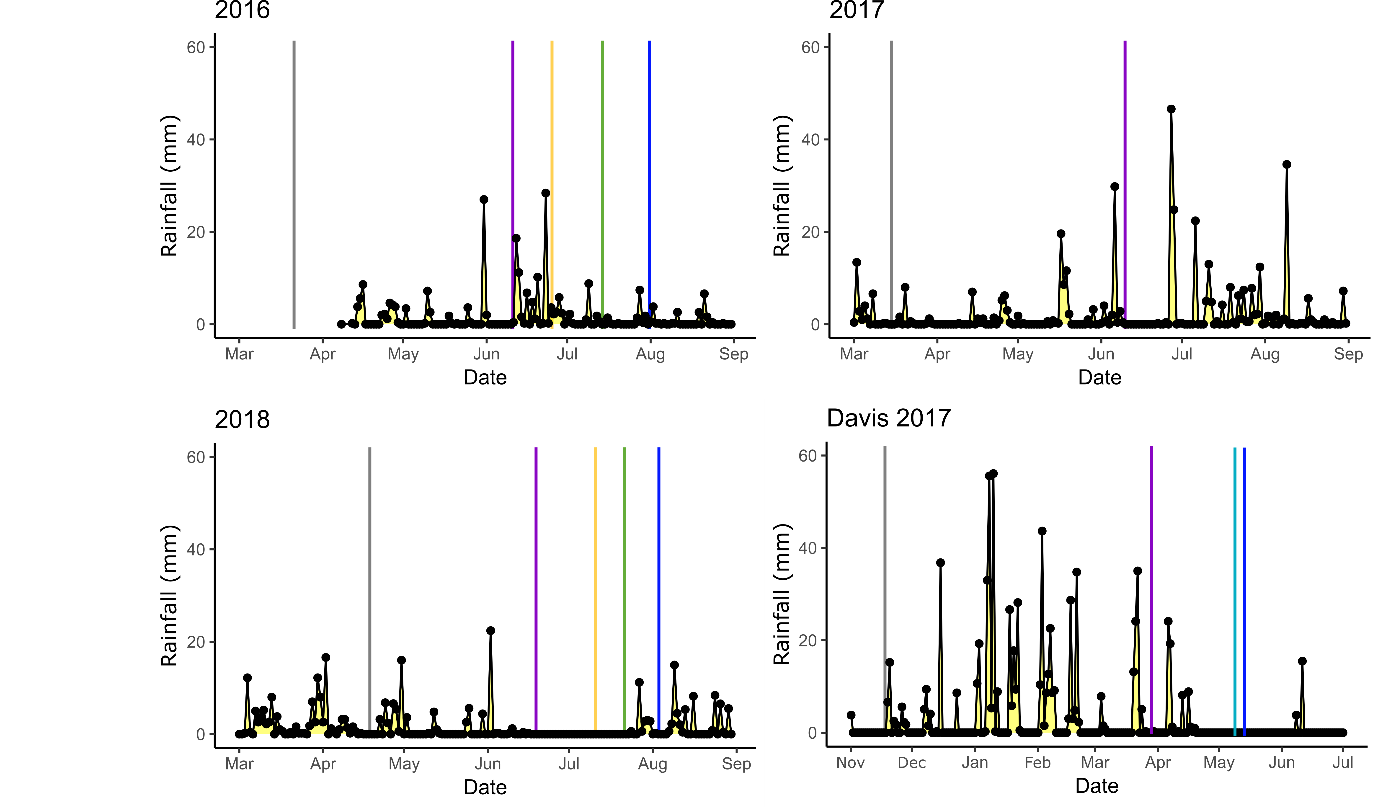

**Supplementary Figure 13: Rainfall patterns for field trials at the JIC and Davis, CA.** Total rainfall is shown per day for the approximate growing season in each field season. Heading date for each trial is indicated with a purple vertical line, leaf senescence is indicated with a green or yellow line, to show the difference in senescence timings for the green and yellow plants, in the JIC, and with a light blue line in Davis, as no difference was observed. Peduncle senescence timing is indicated with a dark blue line. Only heading date is available for 2017 as senescence onset was not scored for this trial. Rainfall is reported in mm (y-axis). For all years, sowing date is shown as a grey line. Rainfall data for the 2016 JIC field trials is only available from 08/04/2016.

**Supplementary Table 1: Kronos TILLING lines selected from 2015 Field trial.**

| **TILLING line** | **Flag Leaf Senescence** | **Peduncle Senescence** | **Grown in 2016** | **Number of F_2_ Populations** |
| --- | --- | --- | --- | --- |
| K0331 | Early | Early | No | N/A |
| K0344 | Normal | Early | Yes | 1 |
| K0382 | Early | Early | Yes | 2 |
| K0689 | Early | Normal | Yes | 2 |
| K0714 | Late | Late | Yes | 2 |
| K1107 | Late | Late | Yes | 2 |
| K1426 | Early | Early | Yes | 2 |
| K2170 | Early | Early | Yes | 2 |
| K2222 | Late | Late | Yes | 1 |
| K2235 | Early | Early | Yes | 2 |
| K2282 | Early | Late | Yes | 2 |
| K2293 | Late | Late | Yes | 2 |
| K2425 | Late | Late | Yes | 1 |
| K2438 | Early | Late | Yes | 2 |
| K2468 | Late | Late | Yes | 1 |
| K2711 | Late | Late | Yes | 2 |
| K2849 | Early | Late | Yes | 2 |
| K3085 | Late | Late | No | N/A |
| K3117 | Early | Normal | No | N/A |
| K3179 | Late | Late | Yes | 1 |
| K4054 | Late | Late | Yes | 2 |

**Supplementary Table 2:** Total number of reads and mapped reads for each experiment, following removal of PCR duplicates. Average depth of coverage was calculated across the 119.2 Mb target exome space for the exome capture reads (see Krasileva *et al.* 2017). For the Kronos WT reads and the Chromosome 3A reads, average coverage was calculated across the length of Chromosome 3A using 1 Mb genomic windows. Note that depth of coverage was not calculated for the Kronos alignment as the scaffolds are not associated to a chromosome.

| **Sample** | **Total Reads** | **Mapped Reads** | **Percentage of reads mapped (%)** | **Sequencing Depth** | **Experiment** |
| --- | --- | --- | --- | --- | --- |
| K2282-23 Green | 45,224,648 | 44,344,449 | 98% | 47 | Exome Capture |
| K2282-23 Yellow | 41,112,551 | 40,353,815 | 98% | 42 |  |
| K2282-28 Green | 44,755,140 | 43,848,635 | 98% | 46 |  |
| K2282-28 Yellow | 82,272,560 | 80,607,768 | 98% | 85 |  |
| Chromosome 3A | 646,829,708 | 641,183,607 | 99% | N/A | Mapping to Kronos Assembly |
|  | 649,434,109 | 642,304,308 | 99% | 82 | Mapping to RefSeqv1.0 |
| Kronos WT | 1,438,873,905 | 1,208,044,854 | 84% | 30 |  |

**Supplementary Table 3: KASP Markers used in mapping.** Genome specific KASP markers were designed for SNPs derived from the three mapping stages, using the exome capture data and the flow-sorted chromosome 3A data mapped against either the Kronos genome or the RefSeqv1.0 genome. SNP position is given for the RefSeqv1.0 genome and was obtained for the Kronos SNPs using BLAST. The HEX tail (WT) and the FAM tail (Mut) are highlighted in red and blue, respectively.

| **Marker** | **WT** | **Mut** | **Common** | **SNP Position** | **SNP Origin** |
| --- | --- | --- | --- | --- | --- |
| SH266 | gaaggtcggagtcaacggattgacatgagccaacagccatc | gaaggtgaccaagttcatgctgacatgagccaacagccatt | gcaggcaaacaaatcaaaatctat | 627966266 | Original Exome Capture |
| SH467 | gaaggtcggagtcaacggattcttaaacacgttgcagatcagg | gaaggtgaccaagttcatgctcttaaacacgttgcagatcaga | ctccaaatcctcccccaaac | 642038467 | Original Exome Capture |
| SH17119 | gaaggtcggagtcaacggattctggtttcaaggaaattggaag | gaaggtgaccaagttcatgctctggtttcaaggaaattggaaa | gtttctctactgtcccaagt | 644827976 | Kronos Genome |
| SH179 | gaaggtcggagtcaacggattcagtcaagtcaggctcc | gaaggtgaccaagttcatgctcagtcaagtcaggctct | gtgctggaacaaatgaa | 647452179 | Original Exome Capture |
| SH123480 | gaaggtcggagtcaacggattagcatcatatctgggttggtc | gaaggtgaccaagttcatgctagcatcatatctgggttggtt | atgatgaattgaagctattgg | 652471882 | Kronos Genome |
| SH044 | gaaggtcggagtcaacggattcggcgtgccagggctttctc | gaaggtgaccaagttcatgctcggcgtgccagggctttctt | cagcccatatctgggatga | 654781044 | RefSeqv1.1 |
| SH838 | gaaggtcggagtcaacggattgaatcagcttctacagatgg | gaaggtgaccaagttcatgctgaatcagcttctacagatga | gctcctcgcctacgggat | 657616838 | Original Exome Capture |
| SH282 | gaaggtcggagtcaacggattgccttgaggacccatggtgg | gaaggtgaccaagttcatgctgccttgaggacccatggtga | ttggatctcactgtgctggg | 659031282 | RefSeqv1.1 |
| SH59985 | gaaggtcggagtcaacggattctagtggtcgttgtccgc | gaaggtgaccaagttcatgctctagtggtcgttgtccgt | cgtgtggcaagcaagagcg | 659031658 | Kronos Genome |
| SH858 | gaaggtcggagtcaacggatttctatcccggagatcttcctc | gaaggtgaccaagttcatgcttctatcccggagatcttcctt | gtggaagcaccagacataca | 663580858 | RefSeqv1.1 |
| SH567 | gaaggtcggagtcaacggattgtacaacccagaagcgtttaac | gaaggtgaccaagttcatgctgtacaacccagaagcgtttaat | gccaatgcaatctccacgga | 667220567 | RefSeqv1.1 |
| SH969 | gaaggtcggagtcaacggattctggcctgcagaatatagatcac | gaaggtgaccaagttcatgctctggcctgcagaatatagatcat | agcacgtcatcatggcctc | 680354969 | Original Exome Capture |
| SH613 | gaaggtcggagtcaacggattgggaaccccgacatccac | gaaggtgaccaagttcatgctgggaaccccgacatccat | ctgcgcagccagcttgtgcaa | 681464613 | Original Exome Capture |
| SH154 | gaaggtcggagtcaacggattccgccgtgatgctgctcgtg | gaaggtgaccaagttcatgctccgccgtgatgctgctcgta | cgtacaccgggtagatga | 686162154 | Original Exome Capture |
| SH538 | gaaggtcggagtcaacggattgaccagatcaggtcatc | gaaggtgaccaagttcatgctgaccagatcaggtcatt | gccagcgacaagtcc | 714034538 | Original Exome Capture |
| SH256 | gaaggtcggagtcaacggatttcgtggatgctcgacctcc | gaaggtgaccaagttcatgcttcgtggatgctcgacctct | tactttatcaggctcagctcaggga | 731226256 | Original Exome Capture |

**Supplementary Table 4: Rainfall amounts (mm) by key developmental points during the field seasons.**

| **Developmental Stages** | **Field Trial** | **Precipitation (mm)** | **Days** |
| --- | --- | --- | --- |
| Sowing to Heading | JIC 2016 | 104.2 | 70 |
|  | JIC 2017 | 114.4 | 68 |
|  | Davis 2017 | 661.1 | 135 |
|  | JIC 2018 | 88.2 | 64 |
| Heading to Leaf Senescence | JIC 2016 WT | 115.4 | 33 |
|  | JIC 2016 Mut | 87 | 14 |
|  | Davis 2017 | 64.7 | 47 |
|  | JIC 2018 WT | 0 | 32 |
|  | JIC 2018 Mut | 0 | 20 |
| Heading to Peduncle Senescence | JIC 2016 | 128.2 | 50 |
|  | Davis 2017 | 64.7 | 51 |
|  | JIC 2018 | 21 | 43 |
| Sowing to Harvest | JIC 2016 | 221.8 | 137 |
|  | JIC 2017 | 331.4 | 136 |
|  | JIC 2018 | 148.6 | 129 |
|  | Davis 2017 | 744.8 | 212 |

**Supplementary Table 5: SNPs identified against the Kronos Genome.** Each SNP is indicated with its location on the respective Kronos scaffold, the depth of coverage at each SNP, and the type of EMS SNP (G>A or C>T). The SNP is also show in relation to the genes which were identified on the scaffold using BLAST; some scaffolds contain more than one gene, and as a result the SNPs are repeated for each gene. SNPs that were used for KASP marker design are also indicated, as are SNPs that were previously used as KASP markers from the exome capture data (SH179 and S838).

| **Kronos Scaffold** | **Location of SNP** | **Type** | **Depth of Coverage** | **Gene** | **Gene Start** | **Gene End** | **Orientation of Gene** | **SNP relation to gene** | **KASP Marker** |
| --- | --- | --- | --- | --- | --- | --- | --- | --- | --- |
| Triticum_turgidum_Kronos_EIv1.1_scaffold_038440 | 22962 | G>A | 132 | TraesCS3A01G397300 | 38722 | 39595 | + | Upstream |  |
| Triticum_turgidum_Kronos_EIv1.1_scaffold_051420 | 17119 | G>A | 93 | TraesCS3A01G397400 | 50086 | 50739 | - | Downstream | SH17119 |
| Triticum_turgidum_Kronos_EIv1.1_scaffold_051420 | 17119 | G>A | 93 | TraesCS3A01G397600 | 16870 | 17568 | + | In Gene | SH17119 |
| Triticum_turgidum_Kronos_EIv1.1_scaffold_011070 | 133732 | C>T | 98 | TraesCS3A01G399300 | 113656 | 113859 | + | Downstream |  |
| Triticum_turgidum_Kronos_EIv1.1_scaffold_057496 | 12729 | C>T | 100 | TraesCS3A01G401200 | 3654 | 4466 | - | Upstream | SH179 |
| Triticum_turgidum_Kronos_EIv1.1_scaffold_012118 | 44138 | G>A | 28 | TraesCS3A01G401800 | 145669 | 146737 | + | Upstream |  |
| Triticum_turgidum_Kronos_EIv1.1_scaffold_044009 | 3007 | C>T | 48 | TraesCS3A01G402400 | 27036 | 28520 | + | Upstream |  |
| Triticum_turgidum_Kronos_EIv1.1_scaffold_070196 | 25155 | C>T | 76 | TraesCS3A01G402700 | 12487 | 13271 | + | Downstream |  |
| Triticum_turgidum_Kronos_EIv1.1_scaffold_019229 | 128018 | G>A | 45 | TraesCS3A01G403000 | 109526 | 110252 | + | Downstream |  |
| Triticum_turgidum_Kronos_EIv1.1_scaffold_004188 | 141367 | G>A | 28 | TraesCS3A01G404400 | 72793 | 73368 | + | Downstream |  |
| Triticum_turgidum_Kronos_EIv1.1_scaffold_004188 | 196608 | G>A | 32 | TraesCS3A01G404400 | 72793 | 73368 | + | Downstream |  |
| Triticum_turgidum_Kronos_EIv1.1_scaffold_004188 | 269110 | G>A | 64 | TraesCS3A01G404400 | 72793 | 73368 | + | Downstream |  |
| Triticum_turgidum_Kronos_EIv1.1_scaffold_030736 | 46408 | C>T | 19 | TraesCS3A01G406600 | 12110 | 14029 | - | Upstream |  |
| Triticum_turgidum_Kronos_EIv1.1_scaffold_030736 | 86495 | C>T | 37 | TraesCS3A01G406600 | 12110 | 14029 | - | Upstream |  |
| Triticum_turgidum_Kronos_EIv1.1_scaffold_000337 | 3898 | G>A | 43 | TraesCS3A01G407300 | 433422 | 434090 | + | Upstream |  |
| Triticum_turgidum_Kronos_EIv1.1_scaffold_000337 | 65290 | G>A | 121 | TraesCS3A01G407300 | 433422 | 434090 | + | Upstream |  |
| Triticum_turgidum_Kronos_EIv1.1_scaffold_000337 | 123480 | G>A | 182 | TraesCS3A01G407300 | 433422 | 434090 | + | Upstream | SH123480 |
| Triticum_turgidum_Kronos_EIv1.1_scaffold_000337 | 255090 | G>A | 410 | TraesCS3A01G407300 | 433422 | 434090 | + | Upstream |  |
| Triticum_turgidum_Kronos_EIv1.1_scaffold_000337 | 358018 | G>A | 48 | TraesCS3A01G407300 | 433422 | 434090 | + | Upstream |  |
| Triticum_turgidum_Kronos_EIv1.1_scaffold_000337 | 374386 | G>A | 396 | TraesCS3A01G407300 | 433422 | 434090 | + | Upstream |  |
| Triticum_turgidum_Kronos_EIv1.1_scaffold_000337 | 378224 | G>A | 19 | TraesCS3A01G407300 | 433422 | 434090 | + | Upstream |  |
| Triticum_turgidum_Kronos_EIv1.1_scaffold_000337 | 412023 | G>A | 23 | TraesCS3A01G407300 | 433422 | 434090 | + | Upstream |  |
| Triticum_turgidum_Kronos_EIv1.1_scaffold_000337 | 514840 | G>A | 191 | TraesCS3A01G407300 | 433422 | 434090 | + | Downstream |  |
| Triticum_turgidum_Kronos_EIv1.1_scaffold_426139 | 344 | C>T | 15 | TraesCS3A01G409000 | 862 | 1542 | - | Downstream |  |
| Triticum_turgidum_Kronos_EIv1.1_scaffold_064761 | 12835 | G>A | 102 | TraesCS3A01G411800 | 16203 | 17105 | - | Downstream |  |
| Triticum_turgidum_Kronos_EIv1.1_scaffold_047726 | 25045 | C>T | 191 | TraesCS3A01G413900 | 21581 | 24114 | - | Upstream | SH838 |
| Triticum_turgidum_Kronos_EIv1.1_scaffold_047726 | 40197 | C>T | 19 | TraesCS3A01G413900 | 21581 | 24114 | - | Upstream |  |
| Triticum_turgidum_Kronos_EIv1.1_scaffold_047726 | 50838 | C>T | 31 | TraesCS3A01G413900 | 21581 | 24114 | - | Upstream |  |
| Triticum_turgidum_Kronos_EIv1.1_scaffold_016263 | 59609 | C>T | 29 | TraesCS3A01G416800 | 68079 | 69020 | - | Downstream |  |
| Triticum_turgidum_Kronos_EIv1.1_scaffold_016263 | 59985 | C>T | 32 | TraesCS3A01G416800 | 68079 | 69020 | - | Downstream | SH59985 |
| Triticum_turgidum_Kronos_EIv1.1_scaffold_016263 | 59609 | C>T | 29 | TraesCS3A01G416600 | 54461 | 54952 | + | Downstream |  |
| Triticum_turgidum_Kronos_EIv1.1_scaffold_016263 | 59985 | C>T | 32 | TraesCS3A01G416600 | 54461 | 54952 | + | Downstream | SH59985 |
| Triticum_turgidum_Kronos_EIv1.1_scaffold_013527 | 159103 | C>T | 45 | TraesCS3A01G418100 | 18253 | 19646 | + | Downstream |  |
| Triticum_turgidum_Kronos_EIv1.1_scaffold_013527 | 159103 | C>T | 45 | TraesCS3A01G418200 | 20113 | 20469 | + | Downstream |  |

**Supplementary Table 6: SNPs identified against the Chinese Spring RefSeqv1.0.** Each SNP is indicated with its location on chromosome 3A of RefSeqv1.0, the depth of coverage at each SNP, and the type of EMS SNP (G>A or C>T). The nearest gene(s), within 1 Kb of the SNP, are also listed. Where a SNP was within 1 Kb of more than one gene, both genes are listed here. SNPs that were used for KASP markers are indicated.

| **SNP Position** | **Type** | **Quality** | **Depth of Coverage** | **Nearest Gene** | **Gene Start** | **Gene End** | **Orientation of Gene** | **SNP relation to gene** | **KASP Marker** |
| --- | --- | --- | --- | --- | --- | --- | --- | --- | --- |
| 654214125 | C>T | 1742.28 | 53 | TraesCS3A02G409800 | 654214362 | 654216953 | + | Promoter |  |
| 654781044 | C>T | 2525.84 | 74 | TraesCS3A02G410700 | 654781914 | 654785094 | + | Promoter | SH044 |
| 655634840 | G>A | 109.768 | 15 | TraesCS3A02G411700 | 655635825 | 655637330 | - | 3' UTR |  |
| 657616838 | C>T | 6099.96 | 191 | TraesCS3A02G413900 | 657613374 | 657615907 | - | Promoter |  |
| 657616838 | C>T | 6099.96 | 191 | TraesCS3A02G414000 | 657616083 | 657620810 | - | 1st exon | SH838 |
| 659031282 | C>T | 960.391 | 29 | TraesCS3A02G416700 | 659026954 | 659038375 | + | 2nd intron | SH282 |
| 659031658 | C>T | 1058.26 | 32 | TraesCS3A02G416700 | 659026954 | 659038375 | + | 2nd intron | SH59985 |
| 660869609 | C>T | 6383.87 | 218 | TraesCS3A02G419100 | 660864796 | 660872768 | + | 7th intron |  |
| 663580858 | C>T | 5425.45 | 167 | TraesCS3A02G422800 | 663579033 | 663599409 | - | 3rd exon |  |
| 664911627 | C>T | 2943.42 | 92 | TraesCS3A02G423300 | 664912319 | 664912531 | + | Promoter |  |
| 667220567 | C>T | 1980 | 68 | TraesCS3A02G424600 | 667220385 | 667220843 | + | 1st exon | SH567 |
| 667220567 | C>T | 1980 | 68 | TraesCS3A02G424500 | 667219018 | 667220345 | + | Downstream of 3' UTR | SH567 |
| 671647297 | C>T | 2379.2 | 72 | TraesCS3A02G428000 | 671639069 | 671646487 | - | Promoter |  |
| 672089627 | C>T | 6236.04 | 182 | TraesCS3A02G429000 | 672088991 | 672089101 | + | 3' UTR |  |
| 674263850 | C>T | 4796.97 | 138 | TraesCS3A02G432900 | 674255517 | 674279708 | + | 1st intron |  |
| 676234801 | C>T | 2844.75 | 93 | TraesCS3A02G434400 | 676207398 | 676243866 | - | 4th intron |  |
| 680354969 | C>T | 1959.56 | 62 | TraesCS3A02G436500 | 680352382 | 680359377 | - | 4th exon | SH969 |

**Supplementary Table 7. 59 High-confidence genes fall within the YES-1 locus.** Genes are listed in their physical order, from marker SH044 to SH59985. Expression data from the developmental timecourse of cv. Chinese Spring and cv. Azhurnaya were used to identify genes with over 0.5 transcripts per million (TPM) in leaf and shoot tissue (column C). The number of developmental stages in which the gene is expressed above 0.5 TPM in the leaf/shoot is listed in column D; see Figure 4. Whether the gene is expressed in the leaves and shoots across vegetative and reproductive development is listed in column E. Columns F-K list the associated Arabidopsis thaliana and Oryza sativa var. japonica orthologues, alongside their functions and relevant references. Orthologues were obtained using Ensembl BioMart.

*Table is provided as additional Excel file.*

**Supplementary Table 8: TILLING Lines with mutations in a putative Mg2+ transporter,**  **TraesCS3A02G414000, had no phenotype in the field.**

| **Line** | **EMS Mutation** | **Amino Acid Mutation** | **Predicted Consequence** | **Het/Hom** | **Presence of chlorosis?** |
| --- | --- | --- | --- | --- | --- |
| K2282 | G>A | G378R | Missense mutation, SIFT = 0.01 | Hom | Yes |
| K3216 | G>A | NA | Splice donor variant; frameshift and premature stop codon. | Het | No |
| K2091 | G>A | NA | Splice acceptor variant; frameshift and premature stop codon. | Hom | No |
| K2468 | C>T | P187S | Missense mutation, SIFT = 0 | Hom | No |
